# MMD scaffolds ACSL4 and MBOAT7 to promote polyunsaturated phospholipid synthesis and susceptibility to ferroptosis

**DOI:** 10.1101/2022.09.01.506096

**Authors:** Vaishnavi V. Phadnis, Jamie Snider, Victoria Wong, Kyle D. Vaccaro, Tenzin Kunchok, Juliet Allen, Zhong Yao, Betty Geng, Kipp Weiskopf, Igor Stagljar, Whitney S. Henry, Robert A. Weinberg

**Affiliations:** Whitehead Institute for Biomedical Research, Cambridge, MA 02142, USA; Harvard Medical School, Boston, MA 02115, USA; Donnelly Centre, University of Toronto, Ontario, Canada; Dana-Farber Cancer Institute, Boston, MA 02115, USA; Department of Molecular Genetics, Temerty Faculty of Medicine, University of Toronto, Ontario, Canada; Department of Biochemistry, Temerty Faculty of Medicine, University of Toronto, Ontario, Canada; Mediterranean Institute for Life Sciences, Split, Croatia; School of Medicine, University of Split, Croatia; MIT Department of Biology, Cambridge, MA 02142, USA; Ludwig/MIT Center for Molecular Oncology, Cambridge, MA 02142, USA

**Keywords:** ferroptosis, lipid metabolism, arachidonic acid, PUFA, ACSL4, MBOAT7, phosphatidylinositol, scaffold, cancer

## Abstract

Ferroptosis is a form of regulated cell death with roles in degenerative diseases and cancer. Ferroptosis is driven by excessive iron-dependent peroxidation of membrane phospholipids, especially those containing the polyunsaturated fatty acid arachidonic acid. Here, we reveal that an understudied Golgi membrane scaffold protein, MMD, promotes susceptibility to ferroptosis in ovarian and renal carcinoma cells. Upregulation of *MMD* correlates with sensitization to ferroptosis upon monocyte-to-macrophage differentiation. Mechanistically, MMD interacts with ACSL4 and MBOAT7, two enzymes that catalyze consecutive reactions in the biosynthesis of phosphatidylinositol (PI) containing arachidonic acid. MMD increases cellular levels of arachidonoyl-phospholipids and heightens susceptibility to ferroptosis in an ACSL4- and MBOAT7-dependent manner. We propose that MMD potentiates the synthesis of arachidonoyl-PI by bridging ACSL4 with MBOAT7. This molecular mechanism not only clarifies the biochemical underpinnings of ferroptosis susceptibility, with potential therapeutic implications, but also contributes to our understanding of the regulation of cellular lipid metabolism.

**Graphical Abstract:** 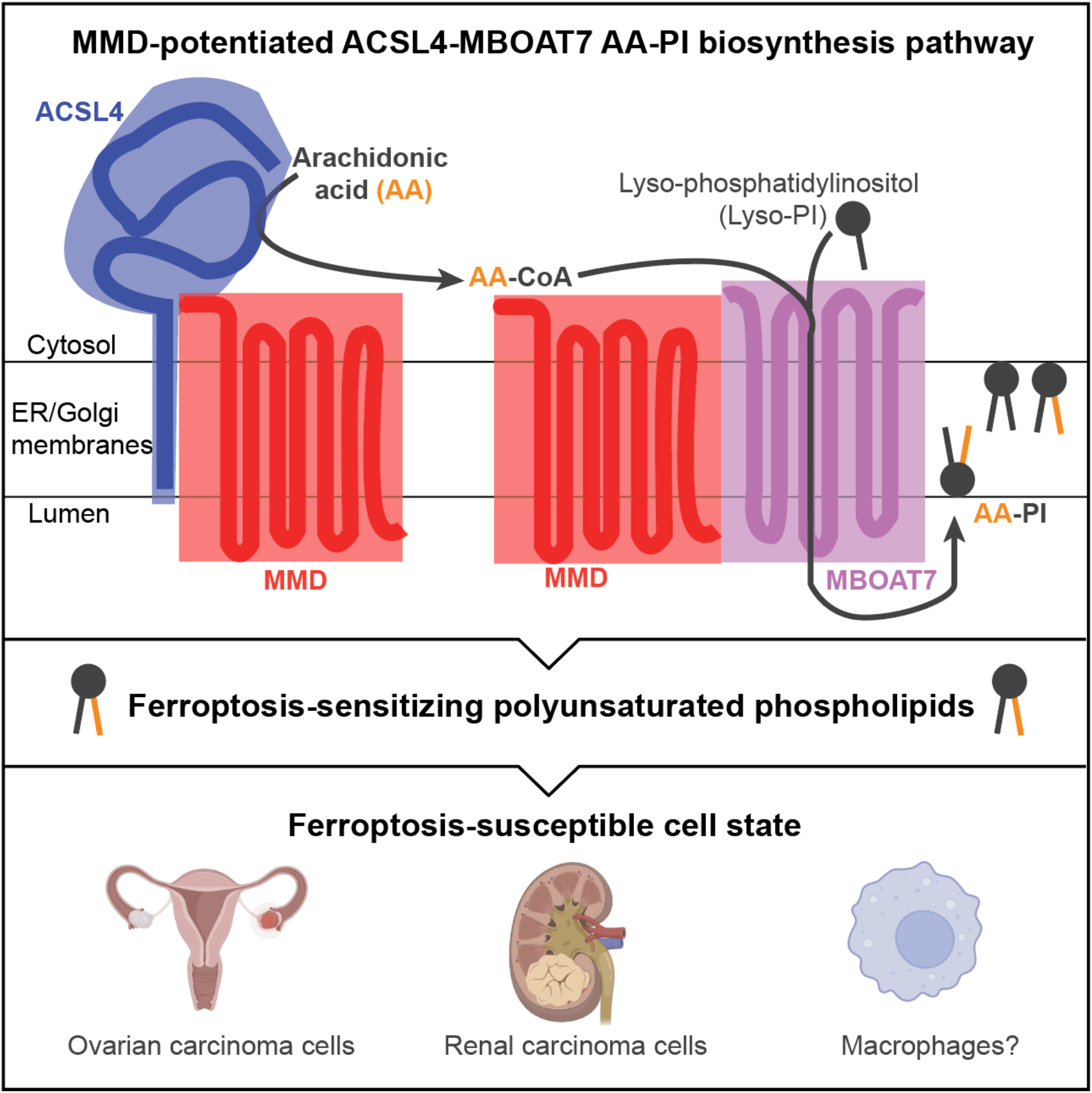

## Introduction

Ferroptosis is a form of non-apoptotic cell death that results from excessive iron-catalyzed peroxidation of membrane phospholipids (Dixon et al., 2012; Jiang et al., 2021). Ferroptosis may contribute to cell death in degenerative diseases and acute injuries of the kidney, liver, heart, and brain (Fang et al., 2019; Guan et al., 2021; Li et al., 2017; Yu et al., 2020). Moreover, ferroptosis inducers exhibit an ability to selectively target aggressive, mesenchymal-enriched and therapy-resistant carcinoma cells (Viswanathan et al., 2017; Hangauer et al., 2017). Mounting evidence suggests that ferroptosis may also be instigated in cancer cells by CD8+ T cells (Wang et al., 2019; Liao et al., 2022). Hence, insights into the molecular mechanisms underlying ferroptosis susceptibility may translate to therapeutic interventions for various pathological conditions, ranging from neurodegeneration to cancer.

Ferroptosis can be induced by pharmacological inhibition of lipid peroxide repair pathways in cells that are poised for runaway lipid peroxidation (Jiang et al., 2021). This primed cell state is characterized by abundant polyunsaturated fatty acid (PUFA)-phospholipids, such as phosphatidylethanolamines (PE) and phosphatidylcholines (PC) containing arachidonic acid (20:4) or adrenic acid (22:4), which serve as critical substrates of peroxidation (Kagan et al., 2017; Zou et al., 2020). Lipid peroxides are generated when bis-allylic hydrogens on the PUFA chains of these phospholipids are abstracted by lipoxygenase enzymes or reactive oxygen species generated by iron-dependent Fenton chemistry (Feng and Stockwell, 2018). Upon inhibition of glutathione peroxidase 4 (GPX4), which normally repairs lipid peroxide damage, a radical chain reaction ensues and leads to ferroptotic cell death (Feng and Stockwell, 2018). Several lipid metabolic enzymes, including ACSL4, LPCAT3, and AGPS, contribute to PUFA-phospholipid synthesis, thereby shaping the ferroptosis-susceptible cell state (Doll et al., 2017; Zou et al., 2020).

We previously reported the findings of two independently performed, genome-wide CRISPR-Cas9 screens in ovarian and renal cancer cells, which revealed genes that promote ferroptosis susceptibility (Zou et al., 2020; Zou et al., 2019). At the intersection of these two screens were 12 candidate genes, the loss of which reduced ferroptosis sensitivity; these included the genes encoding ACSL4, LPCAT3, and AGPS, as well as a gene encoding the poorly studied monocyte-to-macrophage differentiation-related (MMD) protein.

MMD, also termed PAQR11, is a member of the progestin and adipoQ receptor (PAQR) family and functions as an integral endomembrane scaffold protein that promotes vesicle trafficking and mitogenic signaling (Rehli et al., 1995; Tang et al., 2005; Jin et al., 2012; Tan et al., 2017). It was unclear how the known molecular functions of MMD may contribute to increased ferroptosis susceptibility. In addition to its identification in two screens, we noted that *MMD* expression positively correlates with sensitivity to established ferroptosis inducers across cancer cell lines, based on the Cancer Therapeutics Response Portal (Basu et al., 2013; Rees et al., 2016; Seashore-Ludlow et al., 2015). Moreover, *MMD* expression is elevated upon carcinoma cell epithelial-mesenchymal transition (EMT) and promotes metastatic outgrowth, a setting in which ferroptosis induction may offer therapeutic utility (Tan et al., 2021).

Here, we reveal that MMD is a novel regulator of ferroptosis susceptibility in ovarian and renal cancer cells and delineate a molecular mechanism by which it modulates lipid metabolism and ferroptosis susceptibility. We also propose that MMD contributes to dynamic increases in ferroptosis susceptibility in contexts beyond cancer, making it a physiologically relevant modulator of ferroptosis.

## Results

### MMD promotes susceptibility to ferroptosis

MMD was one of only 12 putative pro-ferroptotic genes identified in common between two previously published, independently conducted CRISPR-Cas9 ferroptosis suppressor screens in ovarian and renal carcinoma cells (Figure 1A**, S1A)** (Zou et al., 2020; Zou et al., 2019). Ovarian and renal cancers are both highly aggressive and generally sensitive to ferroptosis inducers, motivating investigation of the role of MMD in the ferroptosis susceptibility of these malignancies (Viswanathan et al., 2017). We also noted that in a survey of 800+ cancer cell lines profiled in the Cancer Therapeutics Response Portal, high expression of *MMD* preferentially correlated with sensitivity to four well-established ferroptosis inducers vs. 400+ other cytotoxic compounds (Figure 1B) (Basu et al., 2013; Rees et al., 2016; Seashore-Ludlow et al., 2015). Given these strong functional and correlative data, we sought to determine whether MMD indeed promotes susceptibility to ferroptosis.

**Figure 1.**
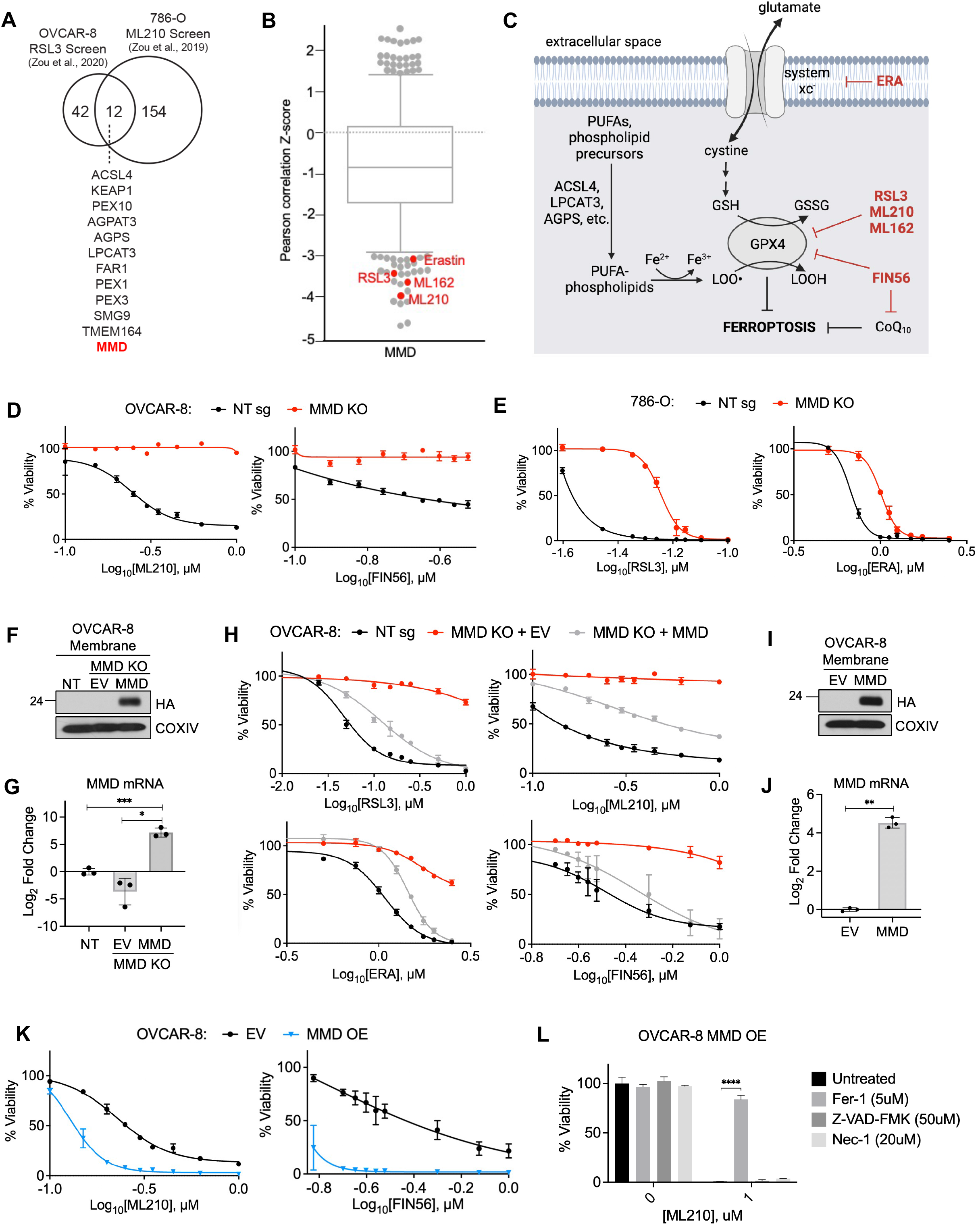
MMD promotes susceptibility to ferroptosis. (See also Figure S1) (A) Venn diagram showing the 12 putative pro-ferroptosis genes identified in two previously published CRISPR screens. Adapted from Zou et al., 2020. (B) Boxplot in which each point represents the Z-scored Pearson correlation coefficient of expression of MMD with area under the viability curve for a distinct cytotoxic drug (481 drugs total), across 860 cancer cell lines. Boxplot interquartile range was set to ∼0.75. Ferroptosis inducers with relatively low Z-scores identified in red. Adapted from the Cancer Therapeutics Response Portal. (C) Schematic of key processes and proteins in ferroptosis, ferroptosis inducing drugs in red. (D) Viability of OVCAR-8 control (NT sg) and MMD KO cells in response to indicated concentrations of ML210 (left) or FIN56 (right). (E) Viability of 786-O control (NT sg) and MMD KO cells in response to indicated concentrations of RSL3 (left) or ERA (right). (F) Immunoblot analysis of membrane proteins in OVCAR-8 NT sg cells, MMD KO cells transduced with empty vector (MMD KO + EV), and MMD KO cells transduced with vector containing HA-MMD (MMD KO + MMD). COXIV was used as a loading control. (G) qRT-PCR analysis of MMD expression in OVCAR-8 MMD KO + EV and MMD KO + MMD cells, compared to NT sg controls. (H) Viability of OVCAR-8 NT sg, MMD KO + EV, and MMD KO + MMD cells in response to indicated concentrations of RSL3 (top left), ML210 (top right), ERA (bottom left), or FIN56 (bottom right). (I) Immunoblot analysis of membrane proteins in OVCAR-8 cells transduced with empty vector (EV) or vector containing HA-MMD (MMD). COXIV was used as a loading control. (J) qRT-PCR analysis of MMD expression in OVCAR-8 MMD OE cells, compared to EV controls. (K) Viability of OVCAR-8 EV and MMD OE cells in response to indicated concentrations of ML210 (left) or FIN56 (right). (L) Viability of OVCAR-8 MMD OE cells in response to ML210 when treated concurrently with Fer-1, Z-VAD-FMK, Nec-1, or nothing (untreated). For all figure panels, data points plotted are mean ± SD of n=3 biological replicates. All experimental figure panels are representative of three independent experiments. ****p < 0.0001, ***p < 0.001, **p < 0.01, *p<0.05

Using CRISPR-Cas9, we established clonal derivatives of the OVCAR-8 (human high-grade serous ovarian carcinoma) and 786-O (human clear cell renal cell carcinoma) cell lines with genetic deletions at the *MMD* locus **(Figure S1B)** and resulting loss of the MMD protein **(Figure S1C)**. Indeed, when compared to control cells expressing a non-targeting control sgRNA (NT sg), MMD knockout (KO) cells were less sensitive to a spectrum of ferroptosis inducers that act via distinct biochemical mechanisms (Figure 1D-E**, S1D-E)**; among these were the covalent GPX4 inhibitors RSL3 and ML210, the cystine import inhibitor erastin (ERA), and the GPX4- and CoQ10-depleting compound FIN56 (Figure 1C) (Yang et al., 2016; Eaton et al., 2020; Dixon et al., 2014; Shimada et al., 2016). Consistent with these observed cell viability responses, MMD KO cells experienced several-fold smaller increases in lipid peroxidation compared to control cells upon ML210 treatment **(Figure S1F-G)**.

To confirm that these effects could be attributed to loss of MMD, we expressed an sgRNA-resistant, HA-tagged MMD cDNA in MMD KO cells (MMD KO + MMD, Figure 1F-G**, S1H)**. Re-expression of MMD partially reverted the sensitivity of MMD KO cells to all four ferroptosis inducers (Figure 1H). Moreover, ectopic overexpression (OE) of HA-MMD in OVCAR-8 parental cells (Figure 1I-J**, S1H)** resulted in hypersensitivity to ferroptosis inducers (Figure 1K**, S1I)**. Further supporting the conclusion that this increased cell death was due specifically to ferroptosis, the viability of ML210-treated MMD OE cells could be rescued by co-treatment with the ferroptosis inhibitor ferrostatin-1 (Fer-1), but not with the apoptosis inhibitor Z-VAD-FMK or necroptosis inhibitor necrostatin-1 (Nec-1) (Figure 1L). Together, these data validated that MMD supports ferroptosis sensitivity in OVCAR-8 and 786-O cells.

*MMD* was originally identified as an mRNA upregulated during monocyte-to-macrophage differentiation (Rehli et al., 1995), and we confirmed this by qRT-PCR in THP-1 monocytes differentiated with phorbol 12-myristate 13-acetate (PMA) **(Figure S1J)** as well as in primary human macrophages vs. monocytes, where we observed 15- to 20-fold upregulation of *MMD* **(Figure S1K)**. To determine whether the role of MMD in ferroptosis may extend to contexts beyond cancer, we investigated whether these increases in *MMD* expression correlate with increased susceptibility to ferroptosis during monocyte-to-macrophage differentiation. Indeed, THP-1 macrophages were more sensitive to ML210 than were THP-1 monocytes **(Figure S1L)**. Similarly, primary human macrophages were far more sensitive to ML210 than monocytes from unmatched human blood donors **(Figure S1M)**. Collectively, these results indicate that MMD is a *bona fide* enabler of ferroptosis susceptibility in ovarian and renal cancer cells, and suggest that its role in ferroptosis may be generalizable to other cancer cell lines and non-neoplastic cells.

### MMD interacts with ACSL4 and MBOAT7 at the endoplasmic reticulum and/or Golgi apparatus

We next pursued the mechanism by which MMD supports ferroptosis susceptibility. MMD is a poorly studied Golgi-resident scaffold protein with no known enzymatic activity (Tan et al., 2017), and its known molecular functions had not previously been directly linked to ferroptosis sensitivity. We therefore explored a recently published proximity labeling interaction dataset that identified hundreds of potential interactors of MMD in H1299 human lung carcinoma cells (Tan et al., 2021). Strikingly, at the intersection of this dataset and the list of hits from both CRISPR screens was a single protein: the 79 kDa membrane-embedded isoform of acyl-CoA synthetase long-chain family member 4 (ACSL4), a PUFA-metabolizing enzyme with an established pro-ferroptotic function (Figure 2A) (Doll et al., 2017).

**Figure 2.**
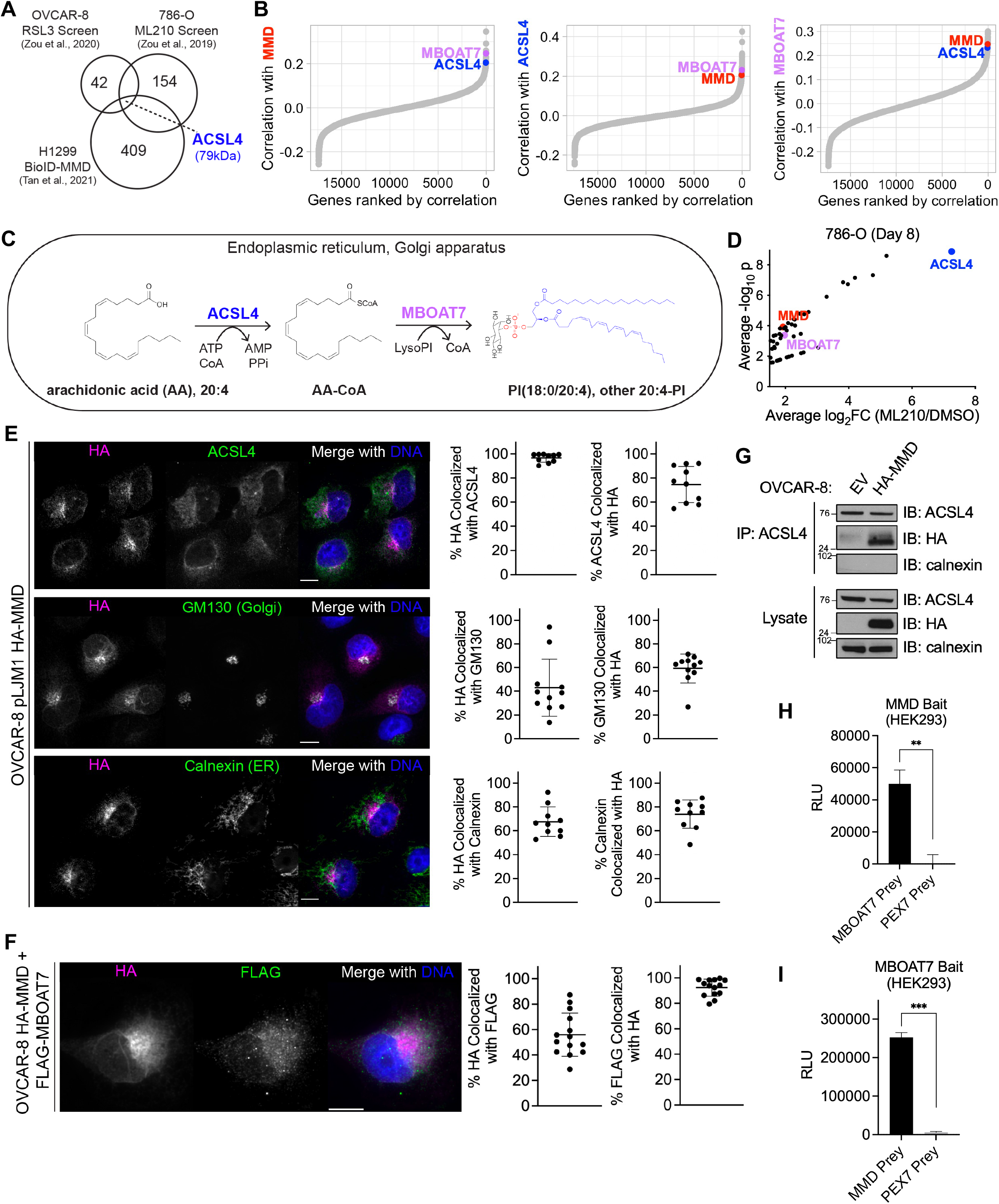
MMD interacts with ACSL4 and MBOAT7 at the endoplasmic reticulum and/or Golgi apparatus. (See also Figure S2) (A) Venn diagram showing the intersection of two previously published CRISPR screens in OVCAR-8 and 786-O cells (Zou et al., 2020; Zou et al., 2019) with a recently published proximity labeling dataset identifying potential interactors of MMD in H1299 cells (Tan et al., 2021). (B) Ordered Pearson correlation coefficients of the effects of loss of MMD (left), ACSL4 (middle), or MBOAT7 (right), with that of loss of every other gene in the genome across all cancer cell lines in DepMap. (C) Schematic showing the consecutive reactions that ACSL4 and MBOAT7 catalyze to generate PI(18:0/20:4). (D) Volcano plot of hits with average log_2_fold change ≥ 1.5 from previously published 786-O CRISPR screen, highlighting MMD, ACSL4, and MBOAT7. Data adapted from Zou et al., 2020. (E) Immunofluorescence staining (left 3 columns) and thresholded Mander’s correlation coefficients (right 2 columns) showing colocalization of HA-MMD with ACSL4 (top row), GM130 (middle row), or Calnexin (bottom row) in OVCAR-8 cells overexpressing HA-MMD. Individual data points are shown, and bars indicate the mean ± SD. Each data point represents the Mander’s correlation coefficient in a distinct field of ∼5 cells on the same coverslip. (F) Immunofluorescence staining (left 3 columns) and thresholded Mander’s correlation coefficients (right 2 columns) showing colocalization of HA-MMD with FLAG-MBOAT7 in OVCAR-8 cells overexpressing both HA-MMD and FLAG-MBOAT7. Individual data points are shown, and bars indicate the mean ± SD. Each data point represents the Mander’s correlation coefficient in a distinct field of ∼5 cells on the same coverslip. (G) Immunoprecipitation and immunoblot analysis of ACSL4 and HA-MMD in OVCAR-8 EV and MMD OE cells. Calnexin was used as a negative control for the immunoprecipitation. (H) Luciferase activity (relative light units, RLU) indicating MaMTH reporter activity in HEK293 cells expressing MMD bait and MBOAT7 prey or PEX7 prey. Data points plotted are mean ± SD of n=3 biological replicates. **p < 0.01 (I) Luciferase activity (relative light units, RLU) indicating MaMTH reporter activity in HEK293 cells expressing MBOAT7 bait and MMD prey or PEX7 prey. Data points plotted are mean ± SD of n=3 biological replicates. ***p < 0.001 All experimental figure panels are representative of three independent experiments.

To further explore whether this potential protein interaction was relevant for MMD’s molecular functions, we mined the DepMap database for codependencies of MMD across hundreds of cancer cell lines (Behan et al., 2019; Meyers et al., 2017). Such codependencies can be used to generate hypotheses about functional similarity and shared membership in a protein complex (Pan et al., 2018). Strikingly, the top 25 codependencies of MMD included not only ACSL4 but also membrane-bound O-acyltransferase family member 7 (MBOAT7) (Figure 2B). MMD, ACSL4, and MBOAT7 mutually ranked highly among each other’s codependencies (Figure 2B), suggesting a functional link.

These observations were noteworthy because ACSL4 and MBOAT7 catalyze sequential reactions to generate certain PUFA-phospholipids from PUFA and lysophospholipid precursors (Figure 2C). Specifically, ACSL4 converts long-chain PUFAs, especially arachidonic acid, to fatty acyl-CoA thioesters that serve as key intermediates in lipid metabolism (Cao et al., 1998; Kuwata and Hara, 2019). MBOAT7, for its part, preferentially esterifies the resulting arachidonoyl-CoA thioesters into lysophosphatidylinositol, leading to the enrichment of arachidonate at the *sn2* position of phosphatidylinositol (PI) (Lee et al., 2008; Caddeo et al., 2021; Xia et al., 2021). While MBOAT7 has not been previously implicated in ferroptosis, MBOAT7 was identified in the 786-O ML210 CRISPR screen (Figure 2D). Accordingly, we hypothesized that MMD physically interacts with both ACSL4 and MBOAT7, juxtaposing two sequentially acting lipid metabolic enzymes to direct arachidonic acid flux toward arachidonoyl-PI synthesis, which in turn may heighten cellular ferroptosis sensitivity.

To test the hypothesis that MMD interacts with ACSL4 and MBOAT7, we first investigated the subcellular localization of MMD via immunofluorescence analyses. Ectopically expressed HA-MMD localized to the Golgi and the endoplasmic reticulum (ER), based on colocalization with both GM130 and calnexin, Golgi- and ER-resident proteins respectively (Figure 2E). Moreover, HA-MMD colocalized with endogenous ACSL4 (Figure 2E) and ectopically expressed FLAG-MBOAT7 (Figure 2F).

Given the spatial congruence of the three proteins, we proceeded to perform co-immunoprecipitation experiments. Corroborating the proximity labeling dataset, we detected HA-MMD in endogenous ACSL4 immunoprecipitates from HA-MMD-expressing cells; conversely, calnexin, a negative control ER protein, was not detected in the ACSL4 immunoprecipitates (Figure 2G). MBOAT7, a large, hydrophobic integral membrane protein, showed very weak signal by immunoblotting even in cells overexpressing FLAG-MBOAT7 (data not shown), and was therefore not conducive to co-immunoprecipitation analysis.

Accordingly, we sought to determine whether MMD could interact with MBOAT7 using the mammalian-membrane two-hybrid (MaMTH) assay (Petschnigg et al., 2014). This assay uses membrane protein “bait” tagged with the C-terminal half of ubiquitin and a transcription factor, and membrane (or soluble) protein “prey” tagged with the N-terminal half of ubiquitin (Petschnigg et al., 2014). Interaction of the bait and prey reconstitutes pseudoubiquitin, which is then cleaved by deubiquitinases. This allows the transcription factor to enter the nucleus and drive transcription of luciferase (Petschnigg et al., 2014). In HEK293 cells, co-expression of MMD bait and MBOAT7 prey (example schematic in **Figure S2A**), or MBOAT7 bait and MMD prey, generated high luciferase activity compared to that of MMD bait or MBOAT7 bait with a negative control peroxisomal protein prey, PEX7 (Figure 2H-I). This evidence serves as proof-of-concept that MMD can interact with MBOAT7 in cells.

Taken together, these data demonstrate that MMD interacts with ACSL4 and MBOAT7. We suggest that MMD simultaneously or sequentially interacts with these consecutively acting enzymes at the ER and/or Golgi, enabling the ACSL4-catalyzed arachidonoyl-CoA intermediate to rapidly encounter MBOAT7 and become incorporated into PI phospholipids.

### MMD increases relative cellular levels of polyunsaturated phospholipids containing ACSL4 and MBOAT7 substrates

The interactions of MMD with ACSL4 and MBOAT7 led us to hypothesize that loss of MMD would lower intracellular levels of certain phospholipid substrates prone to lipid peroxidation during ferroptosis. To pursue this notion, we conducted a mass spectrometry-based, untargeted lipidomic analysis of OVCAR-8 and 786-O MMD KO and control cells. Principal component analysis demonstrated clustering of biological replicates as expected and confirmed the reliability of the dataset **(Figure S3A)**.

For an unbiased assessment of the effects of MMD on the lipidome, we first identified phospholipid species whose relative abundance was statistically significantly altered (p < 0.05) upon loss of MMD in both OVCAR-8 and 786-O cells **(Figure S3B, 3A-C)**. Notably, 15 out of the 17 consistently downregulated species were PUFA-phospholipids, and 11 of these were PUFA-ether phospholipids, all of which are known to increase ferroptosis sensitivity (Kagan et al., 2017; Yang et al., 2016; Zou et al., 2020; Cui et al., 2021) (Figure 3A-C). Moreover, 8 of the 17 downregulated species were similar to oxidized phospholipids categorized as potential ferroptotic cell death signals (see Supplementary Table 1 of Kagan et al., 2017) (Figure 3A-B). Overall, these data indicate that loss of MMD lowers the relative abundance of several ferroptosis-relevant PUFA-phospholipids.

**Figure 3.**
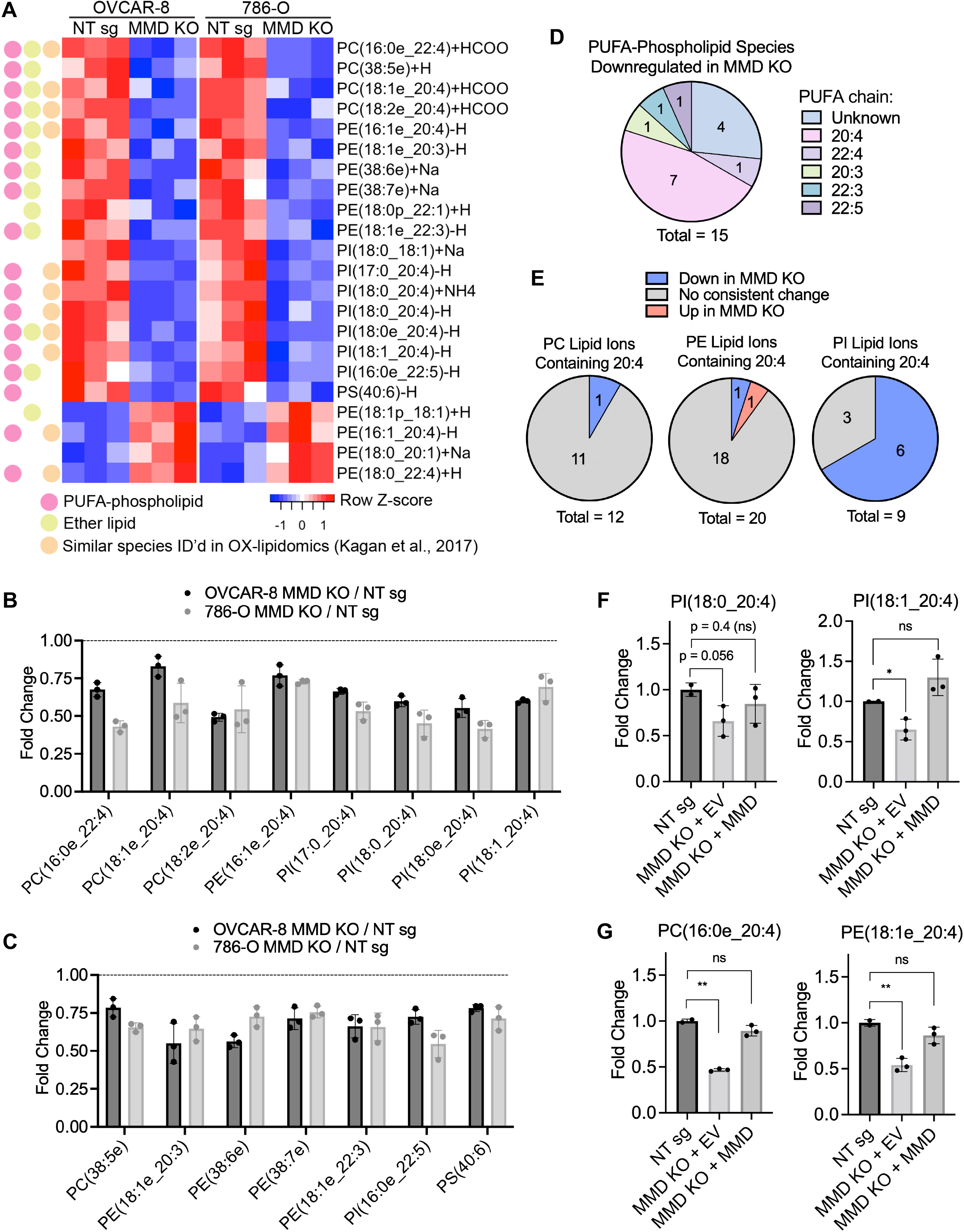
MMD increases relative cellular levels of polyunsaturated phospholipids containing ACSL4 and MBOAT7 substrates. (See also Figure S3) (A) Heatmap showing the 21 lipid species whose relative abundance is consistently (same direction) and significantly (p < 0.05, unadjusted p-value from unpaired, two-tailed, two-sample t-test) altered in both OVCAR-8 and 786-O cells upon MMD KO. Species are annotated as PUFA-phospholipids, ether lipids, and/or ferroptosis-relevant species identified in Kagan et al., 2017. (B) Fold changes of downregulated ferroptosis-relevant species identified in Kagan et al., 2017 from part (A) in OVCAR-8 or 786-O MMD KO cells compared to the respective NT sg cell controls. (C) Fold changes of other downregulated PUFA-phospholipid species from part (A) in OVCAR-8 or 786-O MMD KO cells compared to the respective NT sg cell controls. (D) Distribution of PUFA chains of PUFA-phospholipids that were consistently downregulated in MMD KO cells from part (A). (E) Pie charts, by phospholipid subclass, showing proportions of annotated lipid species containing arachidonic acid which were significantly up- or downregulated in MMD KO cells in part (A). (F) Fold changes of canonical MBOAT7-catalyzed reaction products in OVCAR-8 MMD KO + EV and MMD KO + MMD cells, compared to the NT sg cell controls. Bar plot for NT sg shows the mean ± SD of n=2 biological replicates. (G) Fold changes of other arachidonic acid-containing phospholipids in OVCAR-8 MMD KO + EV and MMD KO + MMD cells, compared to the NT sg cell controls. Bar plot for NT sg shows the mean ± SD of n=2 biological replicates. Unless otherwise specified, data points plotted are mean ± SD of n=3 biological replicates. Panels A-E and F-G are analyses of independent lipidomics experiments. ***p < 0.001, **p < 0.01, *p<0.05

Using the same dataset, we also tested predictions of our model, in which MMD promotes synthesis of arachidonoyl-PI species through ACSL4 and MBOAT7. Among the 11 consistently downregulated PUFA-phospholipids for which the PUFA chain was identified, 7 species contained arachidonic acid (Figure 3D), suggesting enrichment of arachidonate in downregulated phospholipids. We next examined the proportion of arachidonoyl-phospholipids altered upon loss of MMD within each glycerophospholipid class. In line with the preference of MBOAT7 for lysophosphatidylinositol substrates, we noted that 6 of 9 arachidonoyl-PI species were consistently downregulated upon MMD KO, compared to only 1 of 12 arachidonoyl-PC species and 1 of 20 arachidonoyl-PE species detected in our lipidomics experiment (Figure 3E). Given that the major product of MBOAT7-catalyzed reactions is PI(18:0/20:4) (Xia et al., 2021), it is noteworthy that ectopic expression of HA-MMD in MMD KO cells restored the relative abundance of PI(18:0_20:4), PI(18:1_20:4), and PI(18:1e_20:4) (Figure 3F**, S3C)**, confirming that the observed reductions of these species in MMD KO cells were in fact due to loss of MMD. The relative abundances of arachidonate-containing PC and PE species were also restored upon MMD re-expression in MMD KO cells (Figure 3G**, S3D)**, demonstrating that MMD also affects levels of arachidonate-containing phospholipids with headgroups other than PI.

To assess the cell-biological relevance of lower levels of PUFA phospholipids in MMD KO cells, we tested their response to high doses of exogenous palmitate, which preferentially kills cells with elevated saturated-to-unsaturated phospholipid ratios (Piccolis et al., 2019; Zhu et al., 2019). Indeed, MMD KO cells phenocopied ACSL4 KO cells in their increased sensitivity to palmitate-induced cell death **(Figure S3E-F)**, although ACSL4 KO cells were even more sensitive than MMD KO cells, consistent with expected broad reductions in lipid unsaturation upon ACSL4 KO (Zhu et al., 2019). Collectively, these results support the notion that MMD increases levels of ferroptosis-relevant PUFA-phospholipids and drives synthesis of arachidonate-containing PI species through the actions of ACSL4 and MBOAT7.

### MMD drives ferroptosis susceptibility in an ACSL4- and MBOAT7-dependent manner

Our working model, in which MMD bridges ACSL4 and MBOAT7 to promote pro-ferroptotic PUFA-phospholipid remodeling, predicted that MMD would increase ferroptosis sensitivity only in the presence of ACSL4 and MBOAT7. To test this hypothesis, we used CRISPR-Cas9 to deplete either ACSL4 (Figure 4A) or MBOAT7 **(Figure S4A)** in control (EV) and MMD OE cells (Figure 4A, 4C), and then assessed their sensitivity to ferroptosis. Depletion of either ACSL4 or MBOAT7 made control cells resistant to ferroptosis, while MMD OE sensitized control cells to ferroptosis (Figure 4B, 4D). However, MMD OE failed to increase ferroptosis susceptibility in cells lacking either ACSL4 or MBOAT7 (Figure 4B, 4D). This genetic epistasis indicates that MMD requires both ACSL4 and MBOAT7 to increase ferroptosis sensitivity.

**Figure 4.**
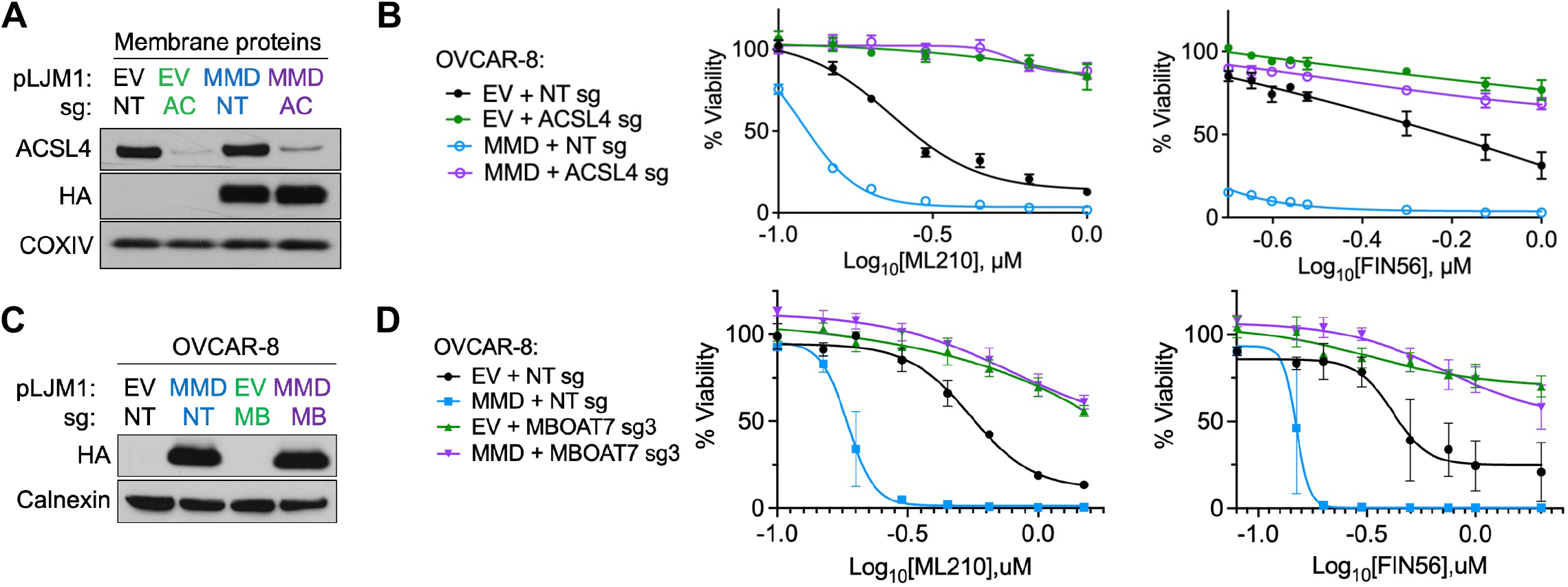
MMD drives ferroptosis susceptibility in an ACSL4- and MBOAT7-dependent manner. (See also Figure S4) (A) Immunoblot analysis of membrane proteins in OVCAR-8 EV cells transduced with a CRISPR vector containing NT sg (EV + NT sg) or ACSL4 sg (EV + ACSL4 sg), and in OVCAR-8 HA-MMD OE cells transduced with a CRISPR vector containing NT sg (MMD + NT sg) or ACSL4 sg (MMD + ACSL4 sg). COXIV was used as a loading control. (B) Viability of cell lines from part (A) in response to indicated concentrations of ML210 (left) or FIN56 (right). (C) Immunoblot analysis in OVCAR-8 EV cells transduced with a CRISPR vector containing NT sg (EV + NT sg) or MBOAT7 sg (EV + MBOAT7 sg), and in OVCAR-8 HA-MMD OE cells transduced with a CRISPR vector containing NT sg (MMD + NT sg) or MBOAT7 sg (MMD + MBOAT7 sg). Calnexin was used as a loading control. (D) Viability of cell lines from part (C) in response to indicated concentrations of ML210 (left) or FIN56 (right). Unless otherwise specified, data points plotted are mean ± SD of n=3 biological replicates, and experimental figure panels are representative of three independent experiments. ****p < 0.0001, **p < 0.01

MBOAT7 is an enzyme within the same family as LPCAT3, the enzyme known to promote ferroptosis downstream of ACSL4 by esterifying PUFA-CoA into PC, PE, and PS lipids (Doll et al., 2017; Hashidate-Yoshida et al., 2015; Zhao et al., 2008). However, unlike MBOAT7, LPCAT3 is not a highly ranked codependency of MMD (data not shown). For the sake of comparison, we performed an epistasis experiment to test whether LPCAT3 may be functionally downstream of MMD as are MBOAT7 and ACSL4. We depleted LPCAT3 **(Figure S4B)** in control and MMD OE cells **(Figure S4C)** and found that, while LPCAT3 depletion alone reduced sensitivity to ferroptosis, MMD OE could sensitize both LPCAT3-WT and LPCAT3-depleted cells to ferroptosis **(Figure S4D)**. Hence, MMD requires ACSL4 and MBOAT7, but not LPCAT3, to promote sensitivity to ferroptosis. Together, these results corroborate the idea that MMD collaborates specifically with ACSL4 and MBOAT7 to render certain carcinoma cells vulnerable to ferroptotic cell death.

## Discussion

Highly aggressive cancer cells exhibiting a mesenchymal gene signature, including drug-tolerant persister cells, EMT-induced cells, sarcomas, and ovarian and renal carcinomas, are difficult to eliminate therapeutically but are often vulnerable to ferroptosis (Hangauer et al., 2017; Viswanathan et al., 2017). In the present study, we undertook to uncover the mechanisms underlying such elevated ferroptosis susceptibility, both to offer insights into the biology of ferroptosis and to guide future efforts to develop ferroptosis-inducing cancer therapies.

We showed that MMD interacts with sequential lipid remodeling enzymes ACSL4 and MBOAT7 and increases the relative cellular abundance of arachidonoyl-phospholipids, thereby heightening ferroptosis susceptibility. Previous reports have suggested that membrane-embedded ACSL3 and ACSL4 promote metabolic flux of arachidonate toward PI via MBOAT7 (Saliakoura et al., 2020; Küch et al., 2014); however, this work provided no mechanistic basis for how the ACSL4-catalyzed intermediate, arachidonoyl-CoA, would preferentially and efficiently encounter MBOAT7 in the crowded intracellular milieu. To explain these previous observations, we propose a mechanistic model in which MMD either simultaneously or consecutively interacts with membrane-embedded ACSL4 and MBOAT7 to enable rapid incorporation of arachidonic acid into PI phospholipids. If this model is further validated, MMD would represent the first integral membrane protein, to our knowledge, that scaffolds metabolic enzymes to increase biosynthetic pathway flux in animal cells (Castellana et al., 2014; Gou et al., 2018).

Our work also reveals multiple new contributors to the ferroptosis-susceptible cell state. LPCAT3 is well-established as an enzyme that promotes ferroptosis sensitivity downstream of ACSL4 (Doll et al., 2017). Our results illustrate that MMD promotes ferroptosis susceptibility through MBOAT7, a protein closely related to LPCAT3 that has not been implicated in ferroptosis. The continued presence of MBOAT7 may explain the relatively modest protective effect of LPCAT3 KO compared to ACSL4 KO that has been observed by some others (Doll et al., 2017). In other words, MBOAT7 may represent an alternative to LPCAT3 in the synthesis of PUFA-phospholipids downstream of ACSL4, with a preference for PI rather than PC, PE, and PS.

The role of PI phospholipids in determining ferroptosis susceptibility warrants further investigation. Intriguingly, a recent report demonstrated that CD8+ T cell-derived IFNγ induces ferroptosis in cancer cells in the presence of exogenous arachidonic acid and does so in an ACSL4-dependent manner (Liao et al., 2022). Upon concurrent IFNγ and arachidonic acid treatment, cancer cells actively incorporated arachidonic acid into many phospholipids, especially PI (Liao et al., 2022). Future work should investigate whether MMD and MBOAT7 are involved in this dynamic remodeling toward arachidonoyl-PI and drive T cell-mediated ferroptosis of cancer cells *in vivo*. If so, activating the MMD-potentiated ACSL4-MBOAT7 axis may enhance tumor cell killing via ferroptosis during checkpoint immunotherapy.

MMD may also be dynamically regulated to increase ferroptosis susceptibility in settings beyond cancer, based on our findings in primary human macrophages and THP-1 cells. Previously published lipidomics data of THP-1 cells showed that arachidonate-containing phospholipids, including PI(18:0/20:4), are significantly upregulated upon PMA differentiation (Zhang et al., 2017), which is consistent with increased MMD levels driving ACSL4-MBOAT7 axis activity and ferroptosis sensitivity upon monocyte-to-macrophage differentiation. Macrophage ferroptosis upon erythrophagocytosis may contribute to atherosclerosis (Liu et al., 2022), making MMD a potential therapeutic target in this context as well.

By regulating the abundance of PI(18:0/20:4), MMD may also affect levels of phosphorylated PI derivatives such as PIP_2_ and PIP_3_, (Shulga et al., 2012), which are critical for signal transduction (Shulga et al., 2012), organelle identity, and membrane fusion (Poccia and Larijani, 2009). Thus, beyond the role of MMD in ferroptosis, the novel molecular function described here may help to explain the pleiotropic actions of this poorly characterized protein, such as its potentiation of Akt and ERK signaling (Huang et al., 2012; Jin et al., 2012; Li and He, 2014; Liu et al., 2012) and vesicle trafficking (Tan et al., 2017).

In summary, our work demonstrates that MMD interacts with key enzymes in arachidonoyl-PI synthesis, clarifying its mechanistic role in ferroptosis and opening many avenues for future investigation in the fields of cell death, cancer biology, signal transduction, and molecular metabolism. Modulating MMD, and in turn the ACSL4-MBOAT7 axis, may also offer therapeutic opportunities to enhance ferroptosis in cancer and inhibit ferroptosis in degenerative diseases.

## Acknowledgments

We thank E.C. Holland at Fred Hutchinson Cancer Research Center, Seattle, WA, for providing laboratory space and support to conduct this research from July 2020 to February 2021 due to circumstances of the COVID-19 pandemic. We are grateful to T. Shibue for providing the pDONR221 plasmid. We thank N. Boehnke for valuable discussions. We acknowledge the following Core Facilities at the Whitehead Institute for their instrumental technical expertise: K. Crowder from the Metabolomics Core, K. Daniels and P. Autissier from the Flow Cytometry Core, and C. Rogers and B. Braswell from the Keck Microscopy Core. We also thank T. Oni, M.G. Vander Heiden, and all members of the Weinberg Lab for helpful discussions.

This work is funded in part by the NIH (P01 CA080111), Breast Cancer Research Foundation, Advanced Medical Research Foundation, Samuel Waxman Cancer Research Foundation and Ludwig Center for Molecular Oncology. Research contributions from the Stagljar lab was supported by funding from the Canadian Cancer Society Research Institute (#703889), Genome Canada via Ontario Genomics (#9427 & #9428), Ontario Research fund (ORF/DIG-501411 & RE08-009), CQDM (Quantum Leap) and Brain Canada (Quantum Leap), and Cancer Research Society (#23235), Genentech Inc. US (CLL-018879), Canadian Institutes for Health Research (CIHR #420989), and Toronto Innovation Acceleration Partners (TIAP L150-066 #72058826). V.V.P. was supported by the MIT Undergraduate Research Opportunities Program, Peter J. Eloranta Summer Research Fellowship, and award number T32GM144273 from the National Institute of General Medical Sciences. K.W. was supported by the Valhalla Foundation, NIH T32 CA09172, A Breath of Hope Lung Foundation, ASCO Conquer Cancer Foundation Young Investigator Award, AACR-AstraZeneca Career Development Award for Physician-Scientists in Honor of José Baselga. W.S.H. was supported by postdoctoral fellowships from the Jane Coffin Childs Memorial Fund and the Ludwig Center at MIT’s Koch Institute. The content is solely the responsibility of the authors and does not necessarily represent the official views of the National Institute of General Medical Sciences or the National Institutes of Health.

## Author Contributions

Conceptualization, V.V.P. and W.S.H.; Methodology, V.V.P., W.S.H., J.S., Z.Y., K.W., and I.S.; Formal Analysis, V.V.P.; Investigation, V.V.P., J.S., V.W., K.D.V., T.K., J.A., Z.Y., and B.G.; Resources, J.S., Z.Y., and I.S.; Writing – Original Draft, V.V.P., W.S.H., and R.A.W.; Writing – Review & Editing, V.V.P., J.S., K.W., W.S.H., and R.A.W.; Supervision, W.S.H. and R.A.W.; Funding Acquisition, K.W., I.S., and R.A.W.

## Declaration of Interests

K.W. declares relationships pertaining to macrophage-directed therapies including patents and royalties (Stanford University, Whitehead Institute, Gilead Sciences); co-founder, SAB member, and equity holder (ALX Oncology, DEM Biopharma); scientific advisor (Carisma Therapeutics). The other authors declare no competing interests.

## Methods

### Cell Line Sources and Culture Conditions

All cells were cultured in a humidified incubator at 37C with 5% CO_2_. OVCAR-8 cells were obtained from J. Brugge (Harvard Medical School) and cultured in 1:1 MCDB 105 medium and Medium 199 supplemented with 10% fetal bovine serum (FBS) and 1% penicillin/streptomycin (P/S). 786-O cells were obtained from the Broad Institute Biological Samples Platform and cultured in RPMI 1640 medium supplemented with 10% FBS and 1% P/S. THP-1 cells were obtained from Jaime Cheah (Koch Institute for Integrative Cancer Research) and cultured in RPMI supplemented with 10% heat-inactivated FBS and 1% P/S. HEK293T cells were obtained from the American Type Culture Collection (ATCC) and were cultured in DMEM supplemented with 10% heat-inactivated FBS. HEK293 cells were obtained from Thermo Fisher (Flp-In 293 TREx cells) and were cultured in DMEM with 10% FBS and 1% P/S. OVCAR-8 and 786-O cells were verified to be free of mycoplasma contamination and STR profiled by the Duke University DNA analysis facility.

### Primary monocyte culture and macrophage differentiation

Discarded blood products were obtained from anonymous human donors and provided by the Crimson Core Specimen Bank (Brigham and Women’s Hospital, Boston, MA). Monocytes were isolated by magnetic labeling with StraightFrom Whole Blood CD14 Microbeads and AutoMacs-directed selection (Miltenyi Biotec). Primary human macrophages were differentiated ex vivo from monocytes by culturing the monocytes in IMDM supplemented with 10% heat-inactivated FBS, 1% P/S, and 20 ng/ml human M-CSF (Peprotech) for 7 days—macrophages derived from this method were sustained in these media conditions and replated as necessary.

### Lentiviral production, transduction of target cells, and antibiotic selection

To produce lentivirus, HEK293T cells were co-transfected with the lentiviral plasmid and VSV-G and psPAX2 packaging plasmids at a 2.1 to 0.2 to 3.7 ratio using polyethylenimine. Lentivirus-containing media was harvested from transfected cells after 48 hours, passed through a 0.45um filter to exclude cells, and added to OVCAR-8 or 786-O cells along with polybrene (10ug/ml). Media was replaced 24 hours after transduction, and selection with puromycin (2ug/ml, OVCAR-8, and 4ug/ml, 786-O) or blasticidin (8ug/ml) began 48 hours after transduction.

### Generation of knockout cell lines

CRISPR-Cas9 lentiviral vectors were generated by cloning sgRNAs into BsmBI-linearized lentiCRISPRv2-puro or lentiCRISPRv2-blast using T4 DNA ligase (sgRNA target sequences in Table 1). OVCAR-8 or 786-O cells were stably transduced with the above lentivirus followed by antibiotic selection as described above. Cells were either maintained as bulk populations (NT sg, EV + NT sg, MMD + NT sg, EV + MBOAT7-sg, MMD + MBOAT7-sg, EV + LPCAT3-sg, MMD + LPCAT3-sg, EV + ACSL4 sg, MMD + ACSL4 sg) or single-cell sorted into 96 well plates to generate single-cell clones (MMD KO and ACSL4 KO cells).

**Table 1.**
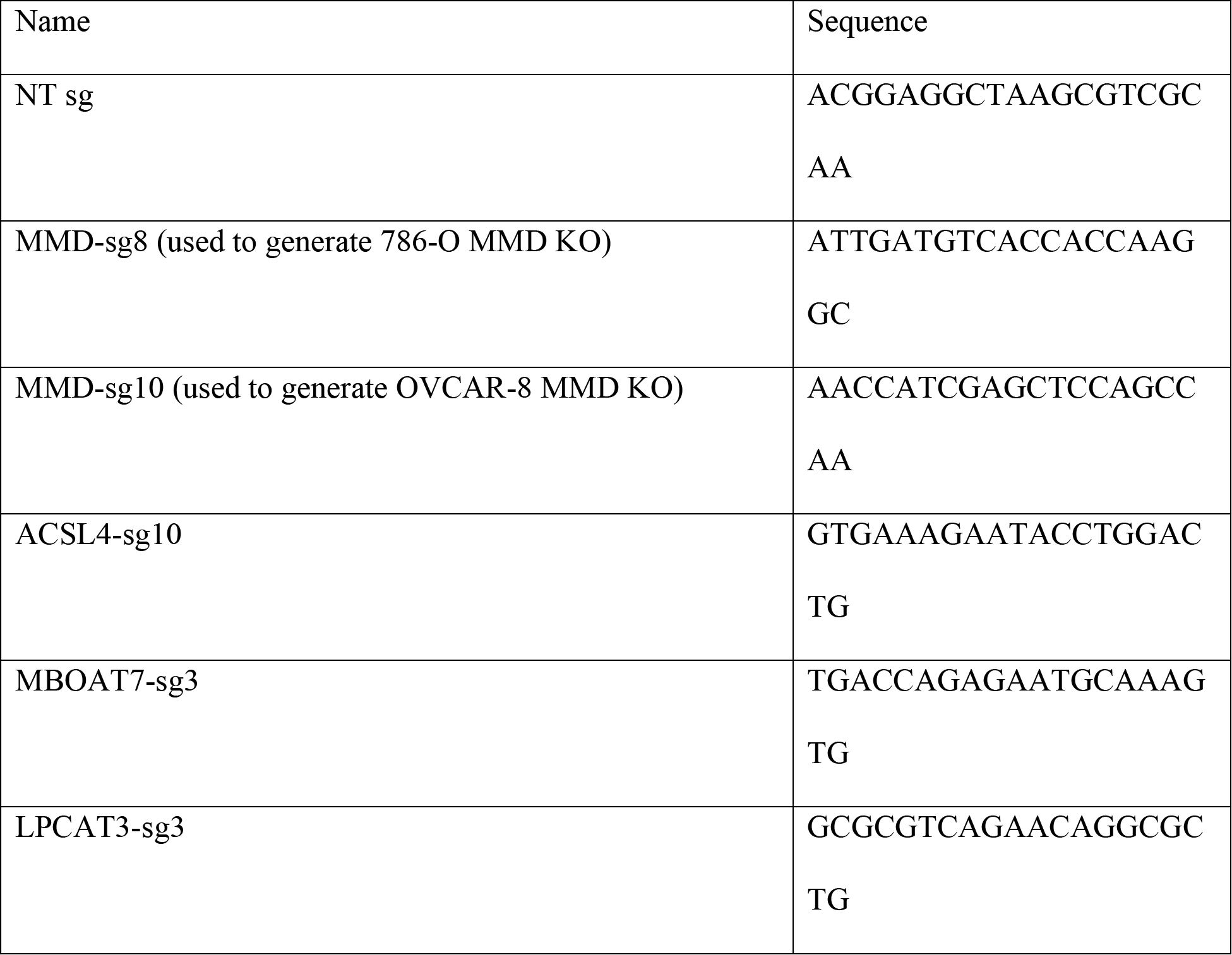
sgRNA target sequences.

### Sanger sequencing validation of MMD knockout cell lines

Single-cell clones generated from bulk MMD-sg cell populations were screened for knockout of MMD at the DNA level. Genomic DNA was purified from cell pellets using the DNeasy Blood & Tissue Kit (QIAGEN). A region of the MMD locus surrounding the sgRNA target site was amplified using PCR and submitted to the Fred Hutchinson Cancer Research Center Genomics Core for Sanger sequencing (all primers in Table 2). Single-cell clones used in this paper were homozygous at the MMD locus. Sequencing reads were aligned to the NCBI MMD reference sequence using ApE.

**Table 2.**
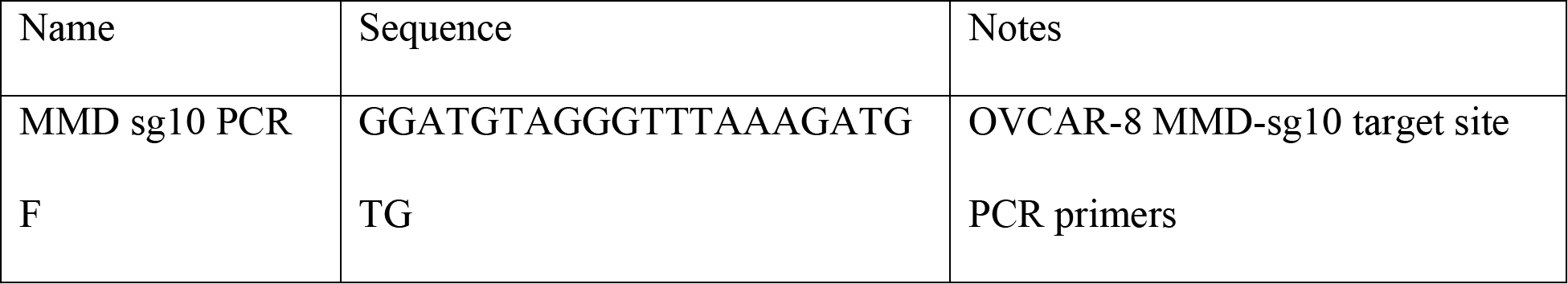
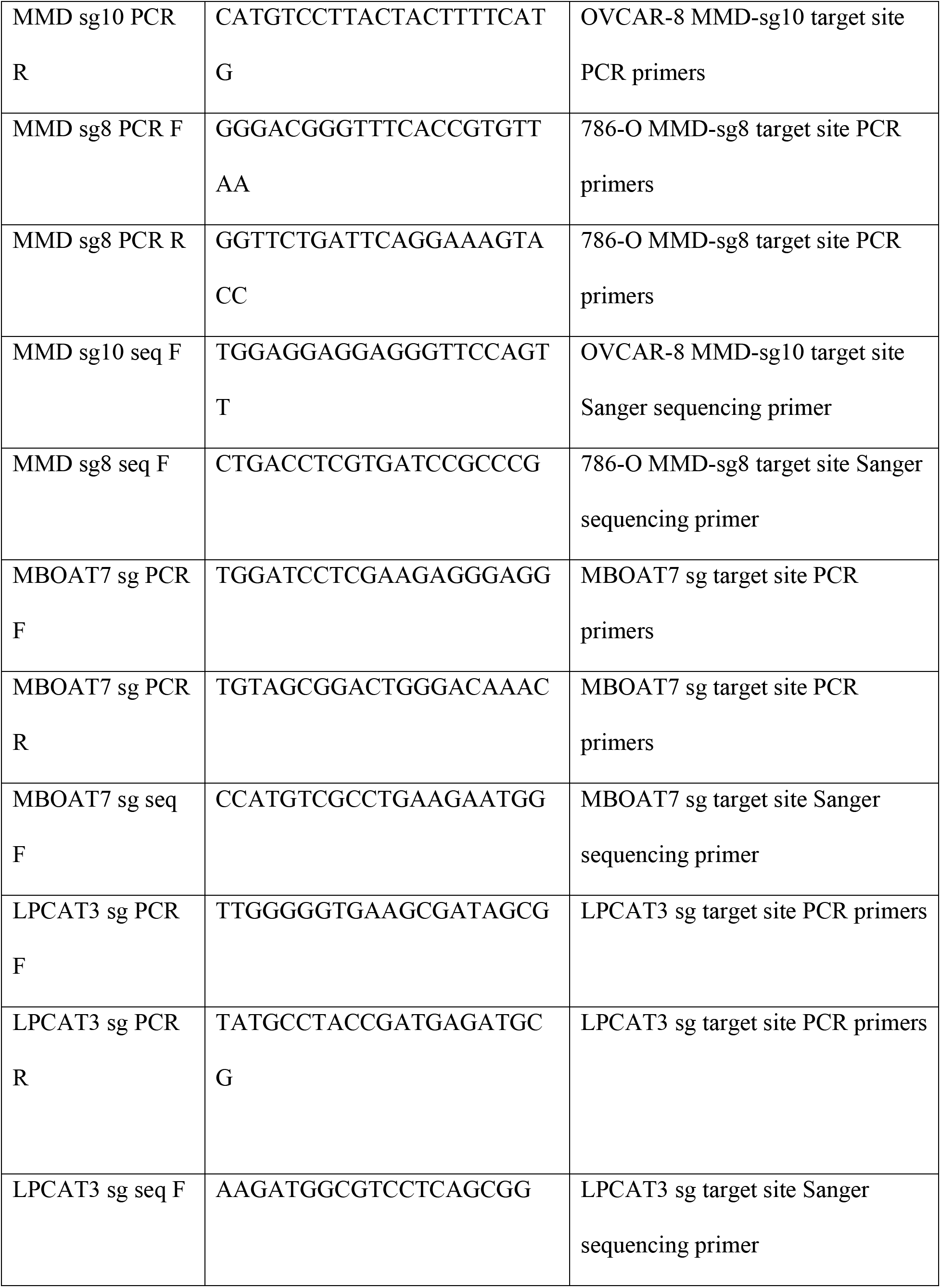
Primers used for Sanger sequencing validation.

### ICE Analysis Validation of MBOAT7 and LPCAT3 depletion

Since high-quality antibodies against MBOAT7 and LPCAT3 were unavailable, MBOAT7 and LPCAT3 depletion in bulk populations of MBOAT7-sg and LPCAT3-sg cells respectively were assessed at the genomic DNA level using Synthego’s Inference of CRISPR Edits (ICE) tool. First, genomic DNA was purified from cell pellets (MBOAT7-sg and LPCAT3-sg as well as the nontargeting control) using the PureLink Genomic DNA Mini Kit (Invitrogen). Regions of the MBOAT7 or LPCAT3 locus surrounding their respective sgRNA target sites were amplified using PCR by Quintara Bio and sequenced using the Sanger method (all primers in Table 2). Subsequently, .ab1 files with mixed peaks in the sg-edited cells were compared to .ab1 files with single peaks in the corresponding control cells within the ICE online tool (https://ice.synthego.com/#/) and the indel score and knockout score were noted.

### Generation of overexpression cell lines

MMD cDNA from the Horizon Discovery Mammalian Gene Collection (BC026324) was fused to an HA-tag at its N-terminus during PCR and cloned into the lentiviral vector pLJM1 linearized with AgeI (NEB) and EcoRI (NEB) by Gibson assembly (NEB). To re-express MMD in MMD knockout cells, the HA-MMD cDNA sequence was edited by fusion PCR (final sequence) to reduce base-pairing with both MMD-sg8 and MMD-sg10 and cloned into the lentiviral vector pLV-EF1a-IRES-blast by restriction digestion with EcoRI-HF (NEB) and BamHI-HF (NEB). FLAG-MBOAT7 was cloned into the lentiviral vector pRH115-mCherry linearized with XbaI (NEB) and BamHI (NEB) by Gibson assembly (NEB) with a codon-optimized FLAG-MBOAT7 gblock (Integrated DNA Technologies). All cDNA inserts were verified by Sanger sequencing (Quintara Bio). Overexpression cell lines were generated by stable transduction of lentiviral vectors and antibiotic selection (pLJM1, pLV-EF1a-IRES-blast) or fluorescence-activated cell sorting for mCherry positive cells (pRH115-mCherry).

### Generation of cell lines for MaMTH experiments

MMD (BC026324) and MBOAT7 (BC003164) cDNAs were obtained from the Horizon Discovery Mammalian Gene Collection and PCR amplified to add attB sites for Gateway cloning (see Table 3). The resulting PCR products were separated on an agarose gel, purified, and combined with the Gateway pDONR221 vector (Invitrogen) in BP Clonase II reactions (Invitrogen) according to manufacturer instructions. The terminated BP clonase reactions were used to transform One Shot™ MAX Efficiency™ DH5α-T1R competent cells (Invitrogen) and transformants were plated on kanamycin agar plates. Single colonies from these plates were verified by Sanger sequencing (Quintara Bio) to contain the desired inserts, and these plasmids were combined with the appropriate destination vectors (see Table 4) in LR Clonase II reactions (Invitrogen) according to manufacturer instructions. The terminated LR clonase reactions were used to transform One Shot™ MAX Efficiency™ DH5α-T1R competent cells (Invitrogen) and transformants were plated on ampicillin agar plates. Plasmids were purified from single colonies and verified to contain the desired inserts (Invitrogen). The protocol used for generation of the MaMTH reporter cell lines was described previously (Saraon et al., 2020).

**Table 3.**
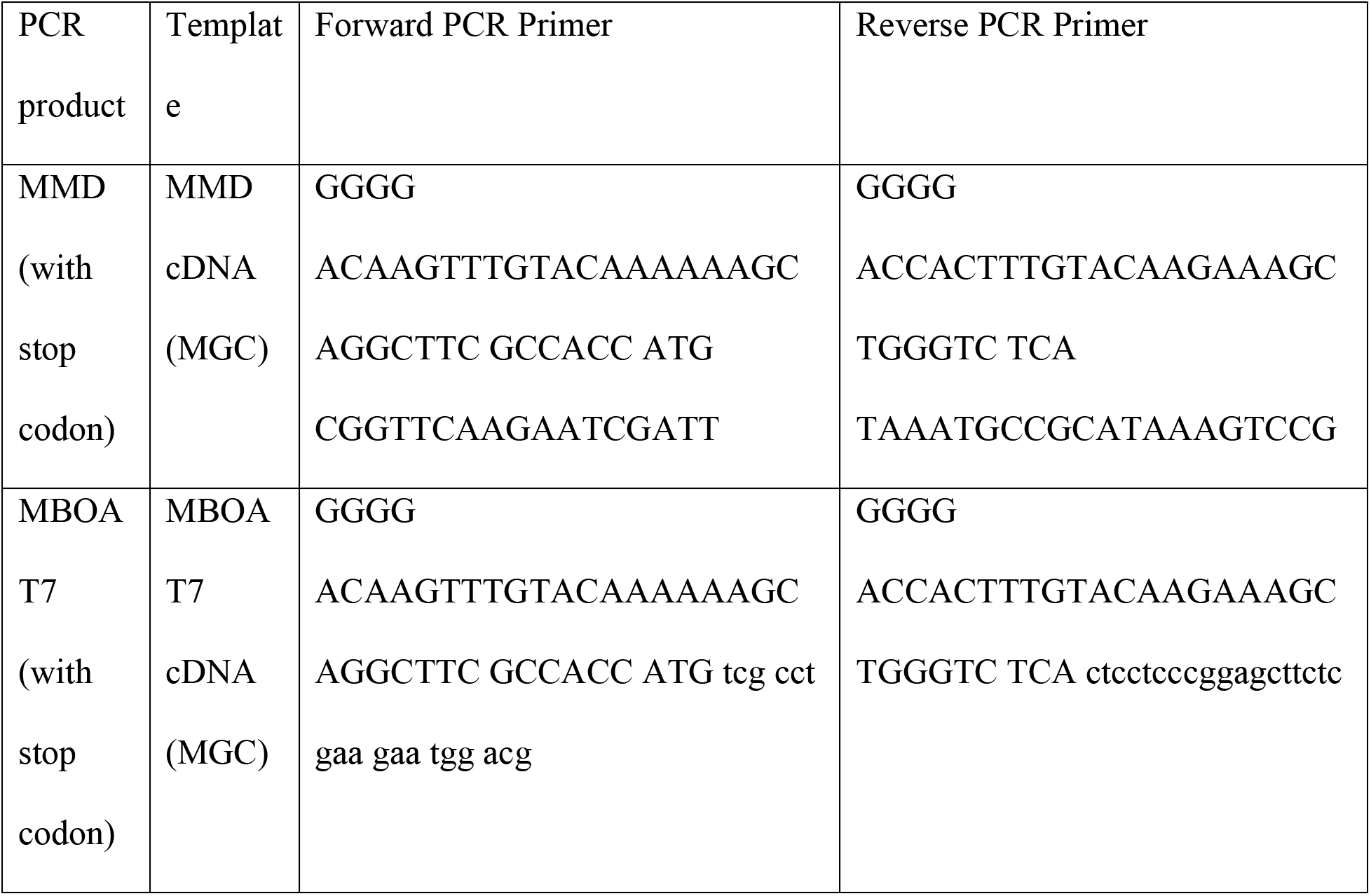
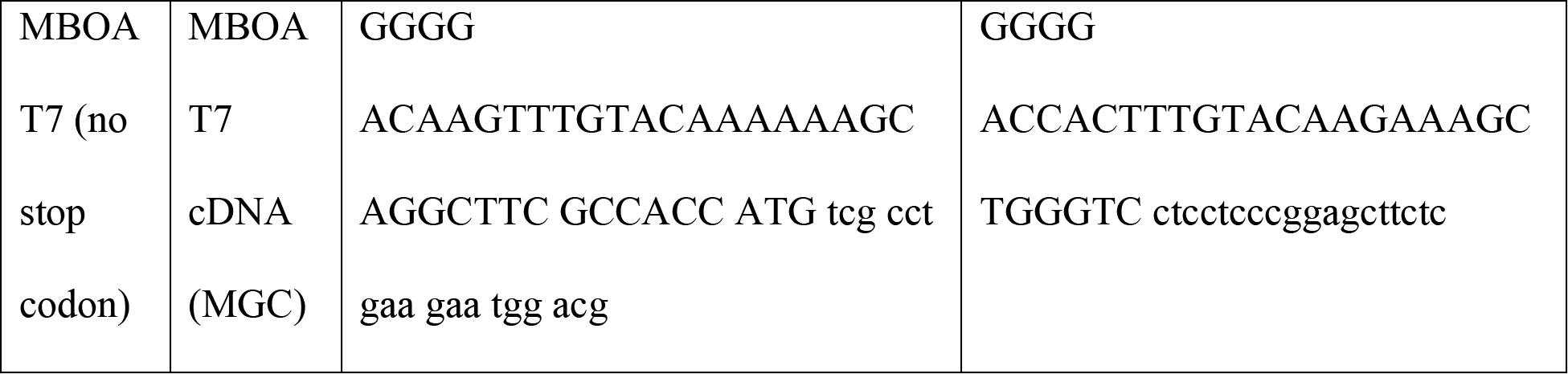
Primers used for addition of attB sites (Gateway cloning)

**Table 4.**
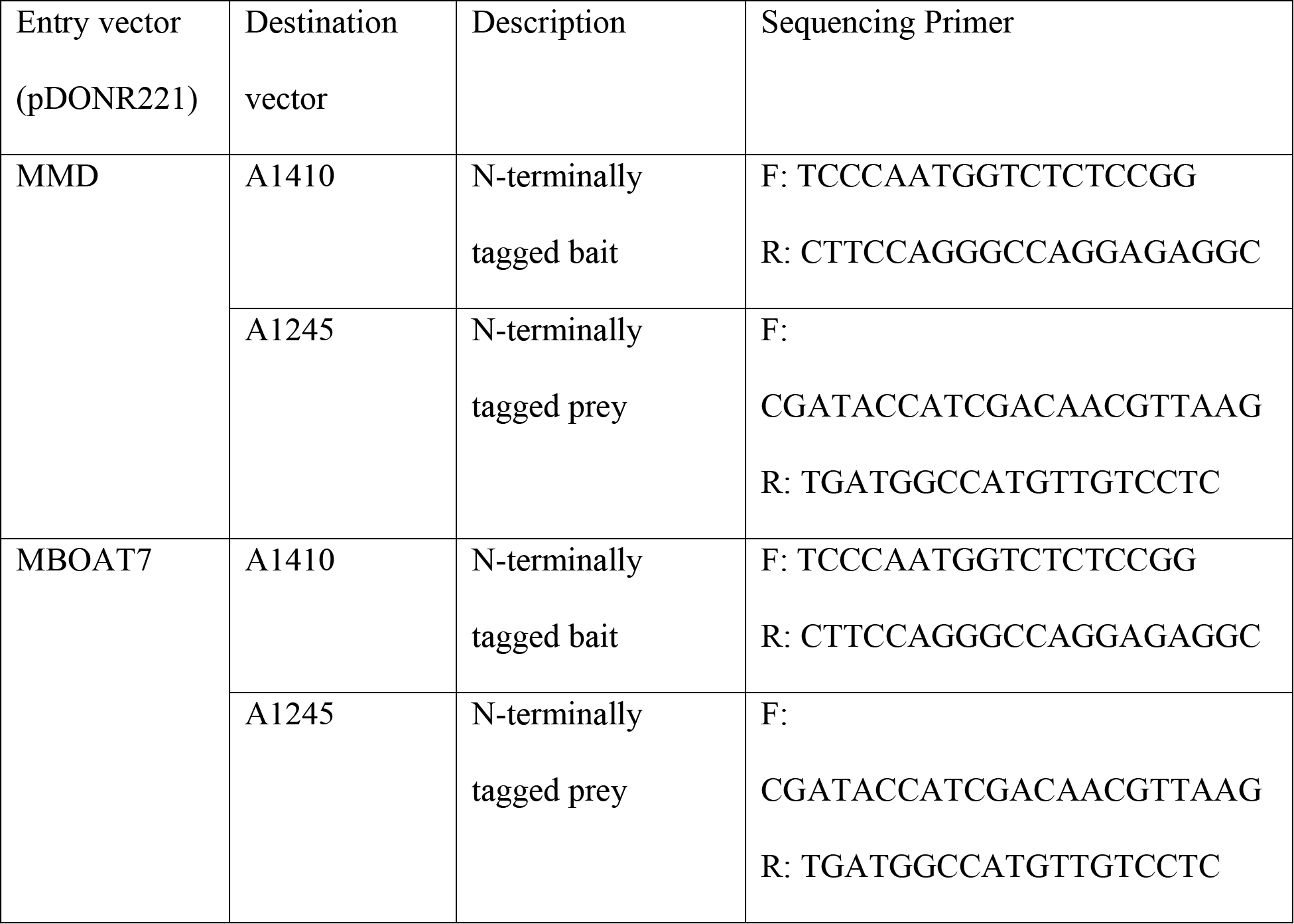
Plasmids for MaMTH experiments.

### Compound Treatment and Cell Viability Assays

For cell viability experiments, cells were treated with a range of concentrations of the indicated small molecule compounds, and relative viability was measured by the CellTiter-Glo Assay. Cells were seeded in 96-well black-wall tissue culture-treated plates at 2,000 to 3,000 cells per well, to reach 30% confluency the following day. Cells were treated with compounds 18-24 hours after seeding using an HP D300e Digital Dispenser with three biological replicates per condition. For BSA-palmitate and BSA-control experiments, cells were treated by manually pipetting. 66-72 hours after compound treatment, viability was measured using the CellTiter-Glo Luminescent Cell Viability Assay (Promega) as per the manufacturer’s instructions. Unless otherwise specified, relative viability was normalized to the untreated condition. Regression fit curves were computed in Prism 8 (GraphPad) using four-parameter inhibition nonlinear regression. The mean and standard deviation for three biological replicates of each data point were calculated. The following compounds were used: RSL3 (Selleck Chem), ML210 (Sigma Aldrich), erastin (Selleck Chem), FIN56 (Selleck Chem), Ferrostatin-1 (Sigma Aldrich), Z-VAD-FMK (Selleck Chem), Necrostatin-1 (Selleck Chem), BSA-palmitate (Cayman Chem), BSA-control (Cayman Chem).

### Lipid Peroxidation Measurement

The redox-sensitive dye, 581/591 C11-BODIPY, was used to measure relative levels of lipid peroxidation. Cells were seeded to 30% confluency and treated with ML210 (1uM for OVCAR-8, 0.25uM for 786-O) for 3.5 hours. 5uM 581/591 C11-BODIPY was also added for the last 30 minutes of the incubation. Cells were trypsinized, washed with PBS 3 times, and strained through a 35um cell strainer for flow cytometry. Cells from each condition were analyzed on a LSRFortessa cytometer using the PE channel for reduced C11-BODIPY and FITC channel for oxidized C11-BODIPY.

### Lysate Preparation

For immunoblots labeled “membrane,” lysates were prepared using the Mem-PER extraction kit using manufacturer buffers and instructions (Thermo). To prepare lysates for other immunoblots, cell pellets were washed in PBS and resuspended in lysis buffer containing 50mM Tris pH 8.0, 150mM NaCl, 1% Triton X-100, 1% digitonin, protease & phosphatase inhibitors (Thermo), and EDTA (Thermo). Lysates were incubated at 4C for 30 minutes with rotation and centrifuged at 4C for 20 minutes at 15,000 rpm in a microcentrifuge. Supernatants were collected and protein concentration was determined by the DC assay using BSA to generate a standard curve.

### Co-immunoprecipitation

For all co-immunoprecipitation experiments, cells from 75-90% confluent plates were washed once with ice-cold PBS and harvested by scraping directly into lysis buffer. Lysates were then prepared and protein was quantified as described above. For each IP, lysate containing 500ug of protein was diluted in 200ul of lysis buffer and pre-cleared by incubating with 10ul of pre-washed Protein A Magnetic Beads (CST #73778) with rotation for 20 minutes at room temperature. IP antibodies were crosslinked to beads by first incubating antibodies with 20ul of pre-washed beads for 30 minutes with rotation at room temperature, then adding 25mM of BS3 crosslinker (Thermo) and rotating for 30 minutes at room temperature, and finally quenching with 100mM Tris (pH 8.0) for 15 minutes with rotation at room temperature. Beads were washed with lysis buffer three times after crosslinking. Cleared lysates were then incubated with antibody-bead conjugates at 4C overnight with rotation. The following day, beads were washed 5 times with lysis buffer and heated in 1X LDS sample buffer with 1X DTT at 70C for 10 minutes. Beads were pelleted and the supernatant was loaded into two lanes as the IP sample for immunoblotting. Generally, 30-40ug of protein was loaded in lysate controls. The following antibodies were used for IP at the indicated concentrations: ACSL4 (Invitrogen PA5-27137, 1:200).

### Immunoblotting Procedure

Samples were prepared by diluting lysates in lysis buffer, NuPAGE LDS Sample Buffer, and NuPAGE Sample Reducing Agent, and heating at 70C for 10 minutes. Samples were resolved by gel electrophoresis using 4-12% Bis-Tris gels and proteins were transferred onto nitrocellulose membranes. Membranes were blocked in 5% milk/TBST, washed 3 times with 1X TBST, and incubated with primary antibodies in 5% BSA/TBST overnight. The following day, membranes were washed with 1X TBST 3 times, incubated with HRP-linked secondary antibodies in 5% milk/TBST for 1 hour, and washed with 1X TBST 3 times. Membranes were then incubated with SuperSignal™ West Dura Extended Duration Substrate (Thermo) or SuperSignal™ West Femto Extended Duration Substrate (Thermo) and developed using X-ray film. The following antibodies were used at indicated concentrations: HA-Tag (rabbit, CST #3724, 1:500 to 1:1000), ACSL4 (rabbit, abcam ab155282, 1:20,000), MBOAT7 (rabbit, Invitrogen PA5-43430, 1:500 to 1:1000), Calnexin (mouse, Invitrogen, MA3-027, 1:1000), COXIV (rabbit, CST #11967, 1:20,000), GAPDH (rabbit, CST #14C10, 1:20,000), HRP-linked anti-rabbit IgG (CST #7074, 1:5000), HRP-linked anti-mouse IgG (CST #7076, 1:5000). The anti-MMD antibody was generated by CST for this work and is currently undergoing commercialization.

### Immunofluorescence Assay

Cells plated on uncoated (786-O) or poly-L-lysine-coated (OVCAR-8) glass coverslips were washed briefly with 1X PBS and then fixed with 4% paraformaldehyde in PBS at room temperature for 10 minutes. After two PBS washes, cells were permeabilized with 0.1% Triton X-100 in PBS at room temperature for 5 minutes. After three more PBS washes, coverslips were blocked in 3% normal donkey serum (NDS) in PBS at room temperature for 1 hour and incubated with primary antibody in 3% NDS/PBS at 4C overnight. For experiments where a fluorophore-conjugated primary antibody was used, the overnight incubation and all following steps were performed in the dark. The next day, coverslips were washed with PBS at room temperature 3 times for 15 minutes each, and then incubated with fluorophore-conjugated secondary antibodies in 3% NDS/PBS at room temperature for 1 hour in the dark. After three more 15-minute washes with PBS at room temperature, coverslips were incubated for 5 minutes at room temperature with DAPI and phalloidin-iFluor 488 in PBS, washed twice with PBS and once with deionized water, and mounted onto slides using Prolong Gold Antifade Mountant. Coverslips were cured at room temperature in the dark for 16-24 hours before imaging. For immunofluorescence experiments staining FLAG-MBOAT7, this protocol was modified to permeabilize cells with 0.1% saponin in PBS instead of Triton X-100, and 0.1% saponin was included in solutions in all subsequent steps until DAPI addition. The following antibodies and fluorescent dyes were used at the indicated concentrations: HA-Tag Alexa Fluor 647 conjugate (mouse, CST #3444, 1:50), HA-Tag (rabbit, CST #3724, 1:250), ACSL4 (rabbit, Invitrogen PA5-27137, 1:250), Calnexin (mouse, MA3-027, 1:250), GM130 (rabbit, CST #12480, 1:3200), DYKDDDDK (FLAG) Tag (rabbit, CST #14793, 1:500), DAPI (0.1ug/ml), phalloidin-iFluor488 (Abcam ab176753, 1:1000), CF 555 donkey anti-rabbit IgG (H+L) 555 secondary antibody (Biotium #20038, 1:500), CF633 donkey anti-mouse IgG (H+L) secondary antibody (Biotium #20124, 1:500).

### Microscopy and Image Analysis

Coverslips were imaged on a DeltaVision Elite Widefield Imaging System using the 60x objective and deconvolved during post-acquisition processing. Quantification of colocalization was performed in Imaris using deconvoluted imaging files. Within Imaris, phalloidin staining was used to mask images so that colocalization was measured only in regions containing cells. Pixel intensity thresholds for colocalized channels were set using negative controls, such that less than or equal to 2 percent of pixels were above each threshold in coverslips stained with secondary antibody only. Mander’s thresholded coefficients in the entire colocalized volume were converted to percentages and presented in figures. Representative images shown in figures were maximally projected and processed in ImageJ.

### MaMTH Assay

MaMTH reporter cells stably expressing *Gaussia princeps* luciferase (New England Biolabs) under the control of a 5xGAL4 UAS were seeded in 96-well tissue culture-treated plates (∼15 000 cells/well) and grown at 37°C/5% CO_2_ overnight in DMEM/10%FBS/1% PS to approximately 50% confluency. Cells were then transiently transfected with MaMTH bait (MMD or MBOAT7) and prey (MMD, MBOAT7 or PEX7 negative control) plasmids using X-tremeGENE9 DNA transfection reagent (Roche) following manufacturer’s instructions. Four hours post-transfection, tetracycline was added to wells to a final concentration of 0.5 µg/mL to induce bait and prey expression. After 48 hours additional growth, luciferase activity was assayed from cell supernatants using 4 µM coelenterazine substrate (Nanolight) and measurement of chemiluminescence with a CLARIOstar plate reader (BMG).

### RNA Purification and RT-qPCR

Cells were pelleted and washed once with PBS. Total RNA was isolated using the QIAgen RNeasy Mini Kit and reverse-transcribed into cDNA using the QuantiTect Reverse Transcription Kit. qPCR was performed on cDNA using SYBR Green Master Mix according to manufacturer instructions. Relative mRNA expression was calculated by the ΔΔCT method with ACTB as reference. The primer sequences are as follows, listed from 5’ to 3’: MMD qPCR Custom F: CCGCTACAAGCCAACTTGCTAT, MMD qPCR Custom R: AGTCCCATTCCATAAATCCATG.

### Untargeted Lipidomics

Lipids were isolated using the following chloroform-methanol extraction method. Cells were seeded in 6 well plates, 24 hours prior, to reach 70% confluency at the time of extraction. Cells were washed with 0.9% cold NaCl on wet ice and then transferred to dry ice. 600 μl LC/MS grade methanol containing 500 nM internal standards (Amino Acid Metabolomics Mix; Cambridge Isotope Laboratories, Inc.) was added to each well. Adherent cells were scraped, collected and samples were transferred to wet ice. 300ul of LC/MS grade water was added, followed by 400ul of chloroform containing Splash LipidoMix (as an internal standard for the lipid fraction; Avanti, Cat. #330707), and samples were vortexed for 10 minutes at 4C. Samples were centrifuged at 4C for 10 minutes at 15,000 rpm in a microcentrifuge and the lower lipid-containing layer was collected and dried using a Speedvac.

Lipids were separated on an Ascentis Express C18 2.1 x 150mm 2.7 um column (Sigma-Aldrich) connected to a Vanquish Horizon UPLC system and an ID-X tribrid mass spectrometer (Thermo Fisher Scientific) equipped with a heated electrospray ionization (HESI) probe. External mass calibration was performed using the standard calibration mixture every seven days. Dried lipid extracts were reconstituted in 50 uL 65:30:5 acetonitrile: isopropanol: water (v/v/v). Typically, 2 uL of sample were injected onto the column, with separate injections for positive and negative ionization modes. Mobile phase A in the chromatographic method consisted of 60:40 water: acetonitrile with 10 mM ammonium formate and 0.1% formic acid, and mobile phase B consisted of 90:10 isopropanol: acetonitrile, with 10 mM ammonium formate and 0.1% formic acid. The chromatographic gradient was adapted from previous work (Bird et al., 2011; Hu et al., 2008). Briefly, the elution was performed with a gradient of 40 min; during 0−1.5 min isocratic elution with 32% B; from 1.5 to 4 min increase to 45% B, from 4 to 5 min increase to 52% B, from 5 to 8 min to 58% B, from 8 to 11 min to 66% B, from 11 to 14 min to 70% B, from 14 to 18 min to 75% B, from 18 to 21 min to 97% B, during 21 to 35 min 97% B is maintained; from 35−35.1 min solvent B was decreased to 32% and then maintained for another 4.9 min for column re-equilibration. The flow rate was set to 0.260 mL/min. The column oven and autosampler were held at 55^∘^C and 15^∘^C, respectively. The mass spectrometer parameters were as follows: The spray voltage was set to 3.25 kV in positive mode and 3.0 kV in negative mode, and the heated capillary and the HESI were held at 300^∘^C and 375^∘^C, respectively. The S-lens RF level was set to 45, and the sheath and auxillary gas were set to 40 and 10 units, respectively. These conditions were held constant for both positive and negative ionization mode acquisitions.

The mass spectrometer was operated in full-scan-ddMS/MS mode with an orbitrap resolution of 120,000 (MS1) and 30,000 (MS/MS). Internal calibration using Easy IC was enabled. Quadrupole isolation was enabled, the AGC target was 1×10^5^, the maximum injection time was 50 msec, and the scan range was *m/z* = 200-2000. For data-dependent MS/MS, the cycle time as 1.5 sec, the isolation window was 1, and an intensity threshold of 1×10^3^ was used. HCD fragmentation was achieved using a step-wise collision energy of 15, 25, and 35 units, and detected in the orbitrap with an AGC target of 5×10^4^ and a maximum injection time of 54 msec. Isotopic exclusion was on, a dynamic exclusion window of 2.5 sec was used, and an exclusion list was generated using a solvent bank.

High-throughput annotation and relative quantification of lipids was performed using LipidSearch v4.2.27 (ThermoFisher Scientific/ Mitsui Knowledge Industries) (Taguchi and Ishikawa, 2010; Yamada et al., 2013) using the HCD database. LipidSearch matches MS/MS data in the experimental data with spectral data in the HCD database. Precursor ion tolerance was set to 5 ppm, product ion tolerance was set to 10 ppm. LipidSearch nomenclature uses underscores to separate the fatty acyl chains to indicate the lack of *sn* positional information (e.g. PC(16:0_18:1) and not (PC(16:0/18:1)). In cases where there is insufficient MS/MS data to identify all acyl chains, only the sum of the chains is displayed (i.e. PC(34:1)). Following the peak search, positive and negative mode data were aligned together where possible (where positive and negative mode data were collected on separate occasions data were aligned separately) and raw peak areas for all annotated lipids were exported to Microsoft Excel and filtered according to the following predetermined quality control criteria: Rej (“Reject” parameter calculated by LipidSearch) equal to 0; PQ (“Peak Quality” parameter calculated by LipidSearch software) greater than 0.75; CV (standard deviation/ mean peak area across triplicate injections of a represented (pooled) biological sample) below 0.4; *R* (linear correlation across a three-point dilution series of the representative (pooled) biological sample) greater than 0.9. Typically, ∼70% of annotated lipids passed all four quality control criteria. Redundant lipid ions (those with identical retention times and multiple adducts) were removed such that only one lipid ion per species/ per unique retention time is reported in merged alignments. For data where positive and negative mode data were aligned separately some redundancies may still exist. Raw peak areas of the filtered lipids were normalized to total lipid signal (positive or negative ionization mode) in each sample to control for sample loading.

Triacylglycerol and cholesteryl ester species were excluded from all further analyses. The measured value of each species in each sample was normalized to the summed values of all lipids in that sample to compare relative abundance of a given lipid species between samples.

### Public Dataset Queries

The Cancer Therapuetics Response Portal was accessed at https://portals.broadinstitute.org/ctrp.v2.1/?page=#ctd2BodyHome on February 23, 2020. Data from the Cancer Dependency Map (DepMap) was accessed at https://depmap.org/portal/ on October 9, 2021.

### Statistical Analysis and Software Information

Statistical analyses were performed in GraphPad Prism and R. Unless otherwise specified, a two-tailed, unpaired student’s two-sample t-test was used to compute p-values. SnapGene and ApE were used for molecular biology analyses. Synthego’s ICE software was used to infer CRISPR editing in bulk populations. Imaris and ImageJ were used for image analyses.

## Supplementary Figure Legends

**Supplementary Figure 1: related to** Figure 1.

(A) Volcano plots of hits with average log_2_fold change ≥ 1.5 from two previously published CRISPR screens, highlighting MMD and ACSL4. Data adapted from Zou et al., 2020.

(B) Alignment of NCBI RefSeq for MMD to PCR-amplified MMD genomic locus in 786-O MMD KO cells (top) or OVCAR-8 MMD KO cells (bottom).

(C) Immunoblot analysis in OVCAR-8 NT sg and MMD KO cells using anti-MMD antibody. GAPDH was used as a loading control. Data are representative of two independent experiments.

(D) Viability of OVCAR-8 NT sg and MMD KO cells in response to indicated concentrations of RSL3 (top) or ERA (bottom).

(E) Viability of 786-O NT sg and MMD KO cells in response to indicated concentrations of ML210 (top) or FIN56 (bottom).

(F) Median fluorescence intensity (MFI) ratio of FITC vs. PE channels in OVCAR-8 NT sg and MMD KO cells treated with redox-sensitive dye C11-BODIPY and DMSO or ML210 (0.5uM). Data show one biological replicate per condition and are representative of three independent experiments.

(G) Median fluorescence intensity (MFI) ratio of FITC vs. PE channels in 786-O NT sg and MMD KO cells treated with redox-sensitive dye C11-BODIPY and DMSO or ML210 (0.25uM). Data show one biological replicate per condition and are representative of three independent experiments.

(H) Immunoblot analysis in OVCAR-8 EV, MMD OE, NT sg, MMD KO + EV, and MMD KO + MMD cells using anti-MMD antibody. SE, short exposure, LE, long exposure. GAPDH was used as a loading control.

(I) Viability of OVCAR-8 EV and MMD OE cells in response to indicated concentrations of RSL3 (top) or ERA (bottom).

(J) qRT-PCR analysis of MMD expression in THP-1 monocytes induced to differentiate into macrophages using PMA, compared to un-induced controls. These data are from one independent experiment.

(K) qRT-PCR analysis of MMD expression in primary human macrophages compared to primary human monocytes from unmatched blood donors. Data points plotted are mean ± SD of n=4 biological replicates.

(L) Viability of THP-1 monocytes (-PMA) and macrophages (+PMA) in response to indicated concentrations of ML210.

(M) Viability of primary human monocytes and primary human macrophages from unmatched blood donors in response to indicated concentrations of ML210. Data points plotted are mean ± SD of n=4 biological replicates.

Unless otherwise specified, data points plotted are mean ± SD of n=3 biological replicates, and experimental figure panels are representative of three independent experiments. ****p < 0.0001, ***p < 0.001, **p < 0.01, *p<0.05

**Supplementary Figure 2: related to** Figure 2.

(A) Example MaMTH setup, corresponding to experiment in Figure 2H. N and C termini of each protein are labeled. Abbreviations: Cub, C-terminal half of ubiquitin; Nub, N-terminal half of ubiquitin; TF, transcription factor, DUBs; deubiquitinase enzymes. When the bait (MMD) and prey (MBOAT7) interact, Cub and Nub are reconstituted as pseudoubiquitin, allowing DUBs to recognize and cleave off the TF. The TF is now free to enter the nucleus and promote expression of the luciferase reporter.

**Supplementary Figure 3: related to** Figure 3.

(A) Principal component analysis of lipidomics dataset.

(B) Schematic of unbiased filtering procedure to identify lipids most likely regulated by MMD.

(C) Fold changes of canonical MBOAT7-catalyzed reaction products in OVCAR-8 MMD KO + EV and MMD KO + MMD cells, compared to the NT sg cell controls. Bar plot for NT sg shows the mean ± SD of n=2 biological replicates.

(D) Fold changes of other arachidonic acid-containing phospholipids in OVCAR-8 MMD KO + EV and MMD KO + MMD cells, compared to the NT sg cell controls. Bar plot for NT sg shows the mean ± SD of n=2 biological replicates.

(E) Immunoblot analysis in OVCAR-8 NT sg and ACSL4 KO cells. GAPDH was used as a loading control. Data are representative of two independent experiments.

(F) Viability of OVCAR-8 NT sg, MMD KO, and ACSL4 KO cells in response to indicated concentrations of BSA-palmitate. Viabilities were normalized to the 750uM BSA control condition. Data are representative of three independent experiments.

Unless otherwise specified, data points plotted are mean ± SD of n=3 biological replicates. Panels A-B vs. C-D are analyses of independent lipidomics experiments. ***p < 0.001, **p < 0.01

**Supplementary Figure 4: related to** Figure 4.

(A) ICE score showing predicted percentage of the bulk population with an insertion or deletion (%indel) or predicted knockout (%ko) at the MBOAT7 locus in the indicated cell lines.

(B) ICE score showing predicted percentage of the bulk population with an insertion or deletion (%indel) or predicted knockout (%ko) at the LPCAT3 locus in the indicated cell lines.

(C) Immunoblot analysis in OVCAR-8 EV cells transduced with a CRISPR vector containing NT sg (EV + NT sg) or LPCAT3 sg (EV + LPCAT3 sg), and in OVCAR-8 HA-MMD OE cells transduced with a CRISPR vector containing NT sg (MMD + NT sg) or LPCAT3 sg (MMD + LPCAT3 sg). Calnexin was used as a loading control.

(D) Viability of cell lines from part (C) in response to indicated concentrations of ML210. Data points plotted are mean ± SD of n=3 biological replicates.

Experimental figure panels are representative of three independent experiments.

