## Supplementary figures and images for "MMD scaffolds ACSL4 and MBOAT7 to promote polyunsaturated phospholipid synthesis and susceptibility to ferroptosis"

### Supplemental Figures

#### Supplementary Figure 1

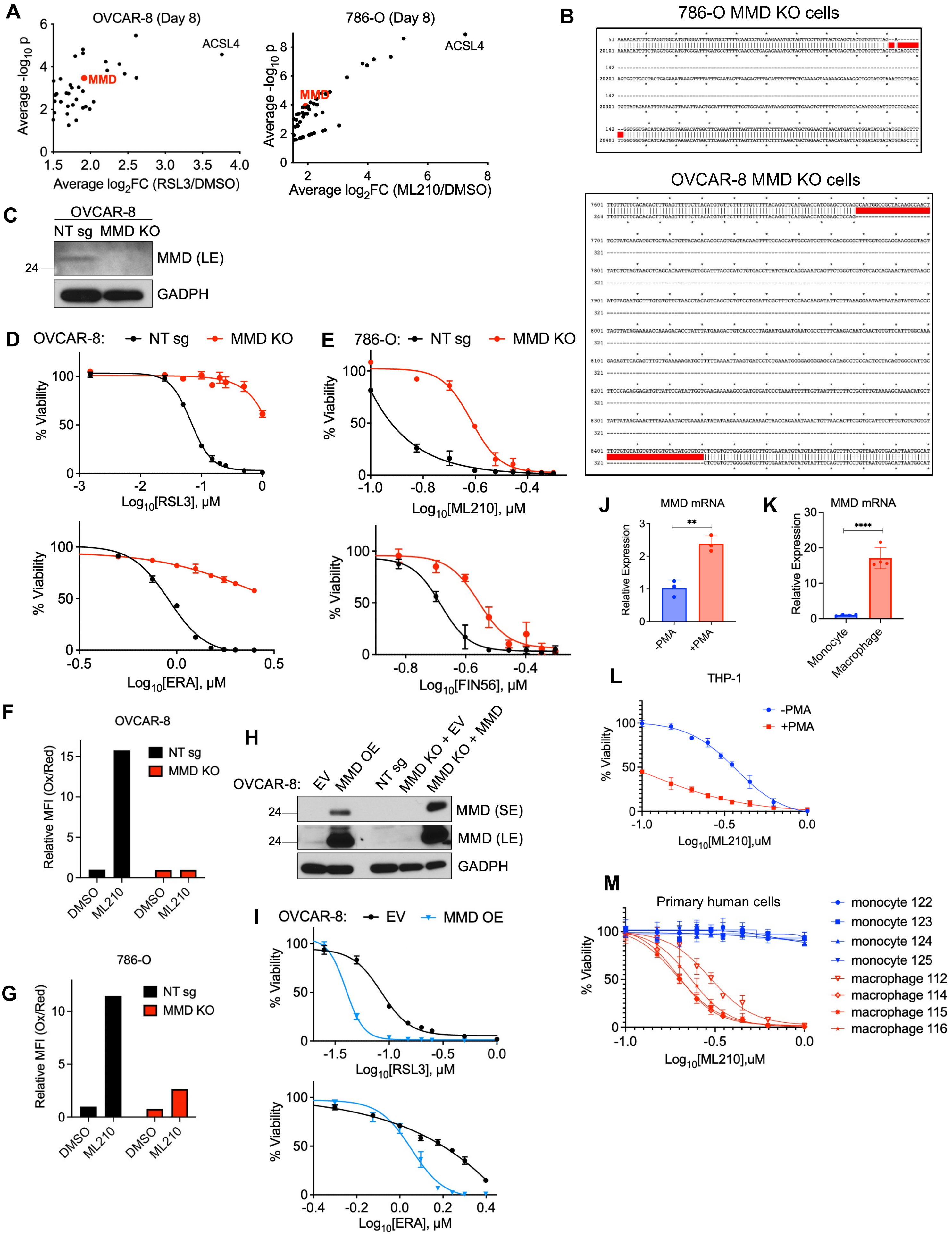

##### Supplementary Figure 2

A

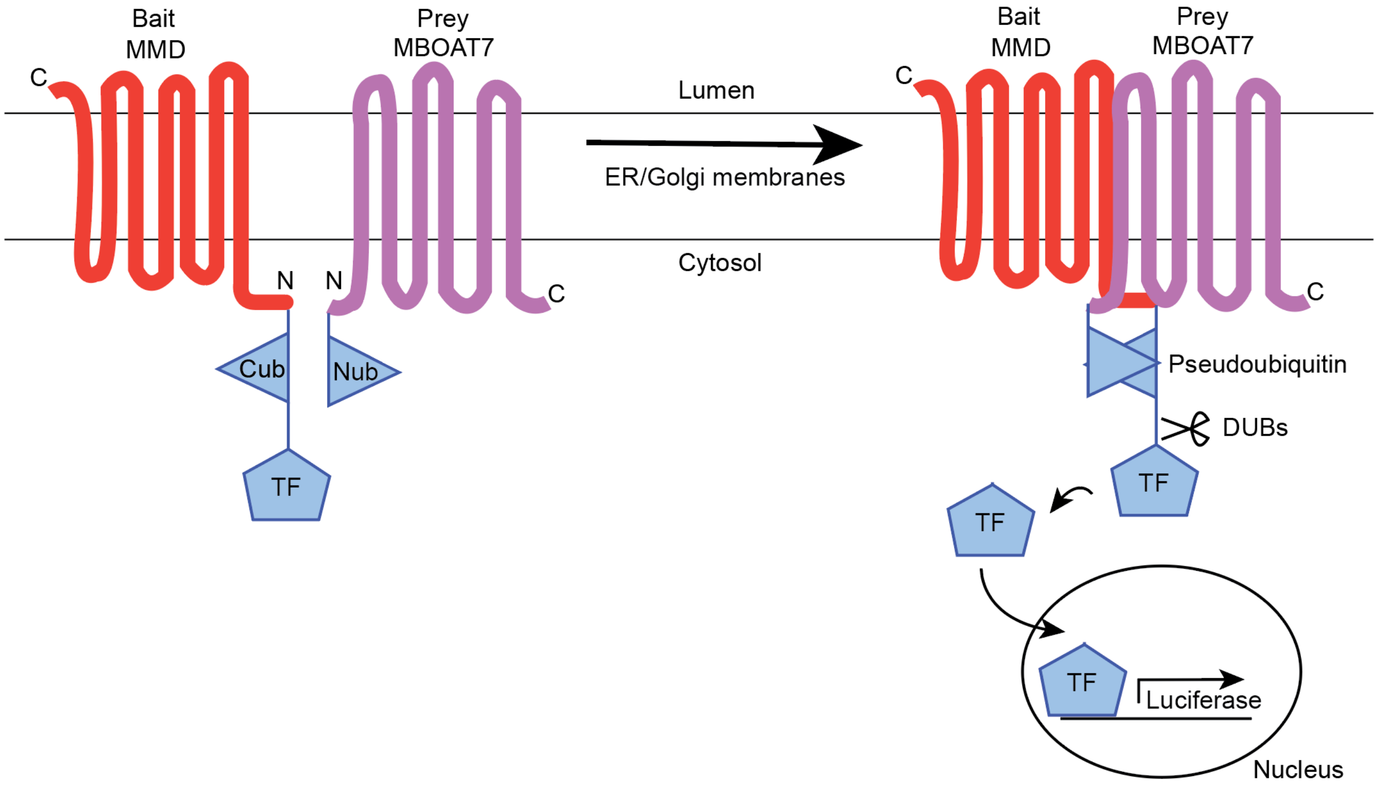

#### Supplementary Figure 3

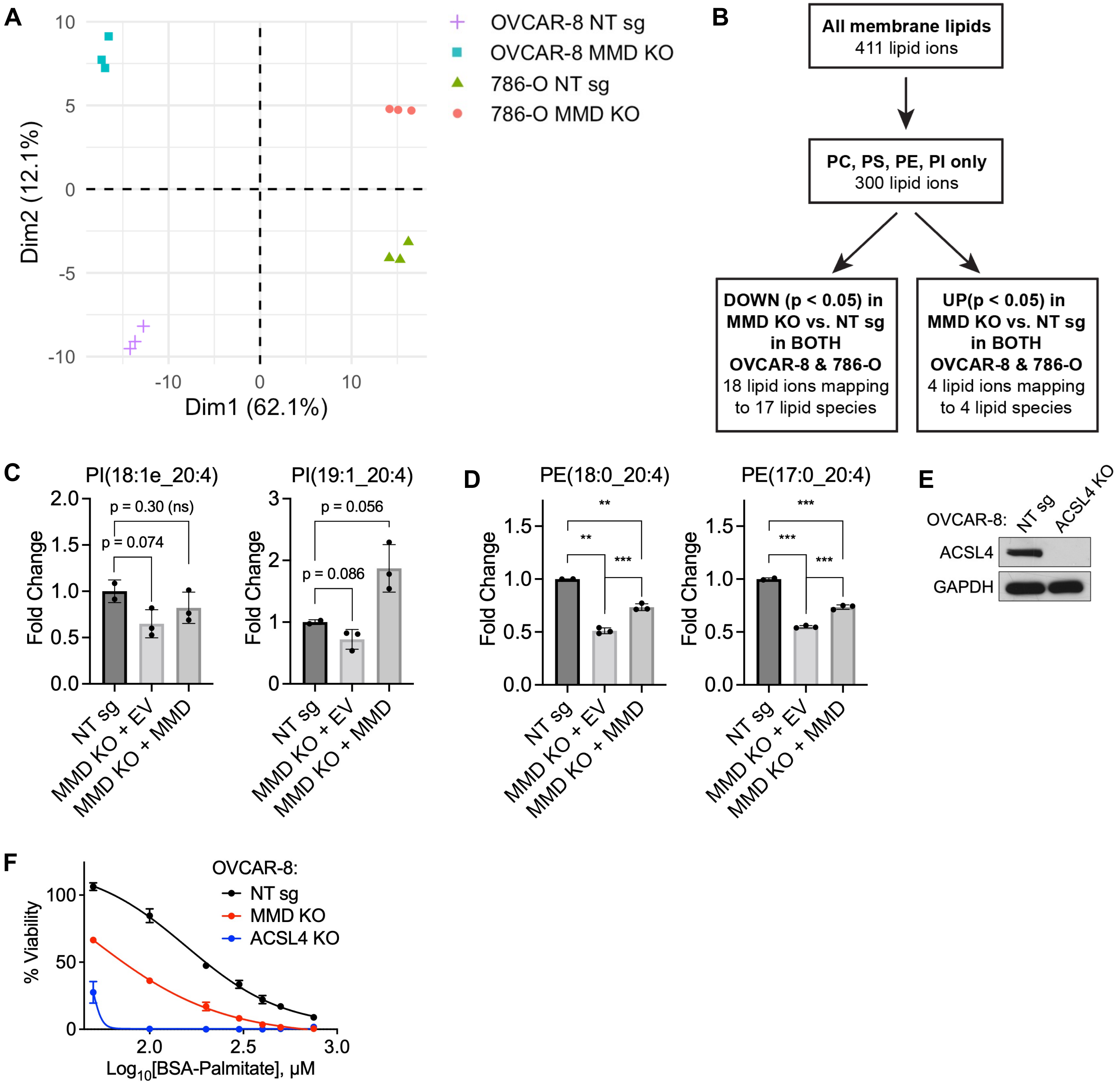

#### Supplementary Figure 4

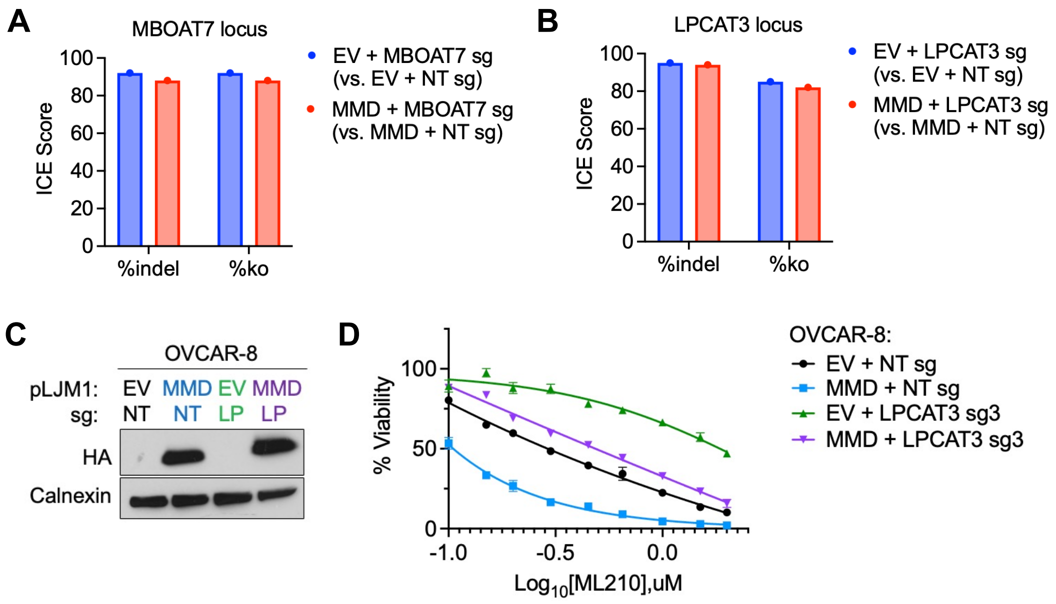
